# Guided by Noise: Correlated Variability Channels Task-Relevant Information in Sensory Neurons

**DOI:** 10.1101/2025.08.13.669902

**Authors:** Ramanujan Srinath, Yunlong Xu, Douglas A. Ruff, Amy M. Ni, Brent Doiron, Marlene R. Cohen

**Affiliations:** Department of Neurobiology and Neuroscience Institute, The University of Chicago, Chicago, IL 60637, USA; Grossman Center for Quantitative Biology and Human Behavior, University of Chicago, Chicago, IL, USA; Department of Statistics, University of Chicago, Chicago, IL, USA

## Abstract

Shared trial-to-trial variability across sensory neurons is reliably reduced when perceptual performance improves, yet this variability is low-dimensional, so it could be ignored by an optimal readout mechanism. Why then is it so consistently related to behavior? We propose that ***shared variability both reflects circuit structure and reveals the information communicated to downstream areas***. In this framework, the same connectivity that shapes signal propagation also shapes shared variability. Using a circuit model, we show that when sensory signals align with shared variability, behaviorally relevant information is amplified without compromising coding fidelity. Analyses of neural population recordings from multiple brain areas and tasks reveal that the dominant axis of *shared variability consistently aligns with task-relevant stimulus features and action plans*. Finally, the behavioral impact of microstimulation can be explained by the extent to which it changes projections onto the shared variability axis. These findings suggest that shared variability may illuminate, rather than obscure, the neural dimensions that guide behavior.

**Significance Statement:** The brain’s ability to use different features of sensory information flexibly across many tasks is essential for complex decision-making. Our study reveals that the correlated variability in neural responses in mid-level visual areas indicates the visual feature that is relevant for behavior. Our biologically plausible network model predicted that it is beneficial for the representation of behaviorally relevant visual information to be aligned with the correlated variability in neurons. We tested and confirmed this in five independent neural data sets. Our results suggest that trial-by-trial variability does not affect the information encoded in sensory neurons but instead is a valuable signal for us to understand which combination of the encoded features is being used by the brain to guide choices.

## Introduction

It is widely reported that correlated neural variability is tightly linked to behavior. Indeed, many studies have shown that good performance on perceptual tasks, associated with factors such as attention, learning, arousal/motivation, or the contrast of a visual stimulus, is consistently accompanied by low shared trial-to-trial response fluctuation in sensory neurons(1–15). Shared variability is often quantified as the mean spike count correlation (also called noise correlations or r_SC_) between the trial-to-trial fluctuations in the responses of pairs of neurons to repeated presentations of the same stimulus(1). Shared variability has become central to experimental and theoretical investigations of information coding, in part because it is modulated by virtually every process known to enhance perception(1, 9, 16), linked to behavior on a trial-by-trial basis(17–20), and sensitive to pharmacological and circuit-level manipulation(5, 21, 22). Further, the structure of shared variability is central to the measured relationships between neurons and behavior(23–25) and impacts the amount of information that is encoded in a neuronal population(16, 26–28). However, the relationships between shared variability, information coding (variability caused by changes in visual inputs), and behavior are complex and have been the subject of ongoing study and debate. Here, we present an overarching framework, supported by data from multiple tasks and brain areas, including both correlative and causal evidence, that can unify many observations.

The many observations and theories in which more shared variability is associated with poorer behavior initially fueled a set of theoretical and experimental investigations about how the nervous system might avoid shared variability in computation. In brief, shared variability interferes with the neuronal population’s ability to encode visual information and, as a consequence, impairs the ability of downstream areas to make accurate perceptual decisions, so it should be avoided(26). From this perspective, cognitive processes such as attention or arousal improve perception by reducing shared noise(29, 30). This should in principle be possible: correlated variability tends to lie within a low-dimensional subspace of neural activity (defined as the space in which the response of each neuron represents one dimension), so that stimulus encoding can be aligned relative to shared variability so as that an optimal decoder can, in principle, ignore it entirely(16, 26–28, 31, 32). Consistent with this, we recently demonstrated that attention dramatically reduces correlated variability in the visual cortex without substantially altering the amount of sensory information that can be gleaned from the population using an optimal linear decoder(33). These observations present a puzzle: if correlated variability need not constrain the sensory information available in a neuronal population, why is it so reliably related to behavior?

Here, we propose a resolution that unifies theoretical and experimental observations: **signal encoding and correlated variability jointly arise from the structure of the cortical circuit, so variability is a reflection of the subspace of neural population activity that guides behavior**. In this framework, while shared variability may not represent the most optimal subspace from an information coding point of view, it provides a window into the information preferentially used to guide behavior. We analyze several datasets to understand the relationship between two types of axes: the axis that accounts for the most shared variability across neurons (correlated variability axis; defined as the dimension that captures the most trial-to-trial variability in population responses in the absence of any visual stimulus), and the axes of signal variability, defined as the dimensions along which task- or stimulus-related responses are encoded. We test the hypothesis that the dimensions of activity that represent the behaviorally relevant visual features are preferentially aligned with the correlated variability axis.

We first test this hypothesis using previously collected V4 population recordings during a change detection task. Detection performance was higher when neuronal responses were better aligned with the correlated variability axis. To understand this relationship, we next studied stimulus estimation in a recurrently coupled network of excitatory and inhibitory neuron models subject to external noise and a separate input signal. In this model, recurrent coupling defines a dominant axis of correlated variability, and signal evoked responses that project onto this axis are selectively amplified by the same circuitry, increasing the information available to a linear decoder.

We evaluated the generality of this principle across multiple datasets by analyzing how correlated variability relates to representations of relevant and irrelevant stimulus features, motor plans, and decisions. First, in a curvature estimation task, when monkeys generalized across irrelevant visual features, the axis of correlated variability captured curvature information across heterogeneous shapes, irrespective of irrelevant features. Second, in the same task, when monkeys instead generalized across variations in saccade target locations, correlated variability more closely reflected motor plans than sensory inputs. Third, in a task where animals alternated discriminating curvature and color, learning to make decisions about a previously irrelevant feature increased the projection of the representation of that feature on the axis of correlated variability. Fourth and finally, we provide causal support for this framework by demonstrating the behavioral impact of microstimulation was predicted by the alignment between evoked population responses and the axis of correlated variability.

Together, these findings challenge the idea that neural representations should, or even possibly could, avoid subspaces of neural population space corrupted by correlated variability. Instead, we suggest that correlated variability arises from the functional structure of the circuit, which reflects the rich landscape of internal states, goals, and sensory and premotor processes that will impact behavior. Correlated variability is preferentially aligned with, rather than orthogonal to, the representations of task-relevant information. Our findings unify and provide a new perspective on decades of experimental results, suggesting that correlated variability may serve as a window into the neural computations that drive perception and decision-making.

## Results

We investigated our central hypothesis by analyzing the relationship between task-relevant information coding in the visual cortex, correlated variability, and behavior. In each data set, we defined the axis of correlated variability as the first principal component (PC1) of neural population activity(34, 35) during a baseline period where rhesus monkeys (*Macaca mulatta*) fixated a central dot on a gray screen at the start of each trial, capturing the dimension that explains the most shared variability in spontaneous activity and used as the reference axis for subsequent analyses. This was largely equivalent to defining the axis based on population responses to repeated presentations to identical stimuli during fixation in our datasets (Figure S1). In all our data sets, the shared variability was low dimensional. The majority of the variance in baseline responses was captured by the first PC, and the second PC accounted for no more than about 10% of the variance (Figure S2). Because of this low dimensional structure, we focus on the first PC when considering how responses align with correlated variability.

### Performance in an orientation change detection task is related to the alignment between the orientation representation and the axis of correlated variability in area V4

We began by testing our hypothesis in a well-established orientation change detection task(2, 9, 12, 20, 33, 35, 36) (Figure 1A). Different aspects of these data have been published previously(21, 35). In this task, monkeys fixated a central dot while two Gabor stimuli of the same orientation were flashed repeatedly. At a random and unsignalled time, one stimulus changed orientation, and the monkey was rewarded for making a saccade to its location. In this task variation, the starting orientations of the gratings varied between 0° and 180° across trials, while the magnitude of the orientation change was fixed. Our analysis focused on trials in which the change occurred at a cued location that overlapped with the joint receptive fields of the recorded V4 neurons.

**Figure 1.**
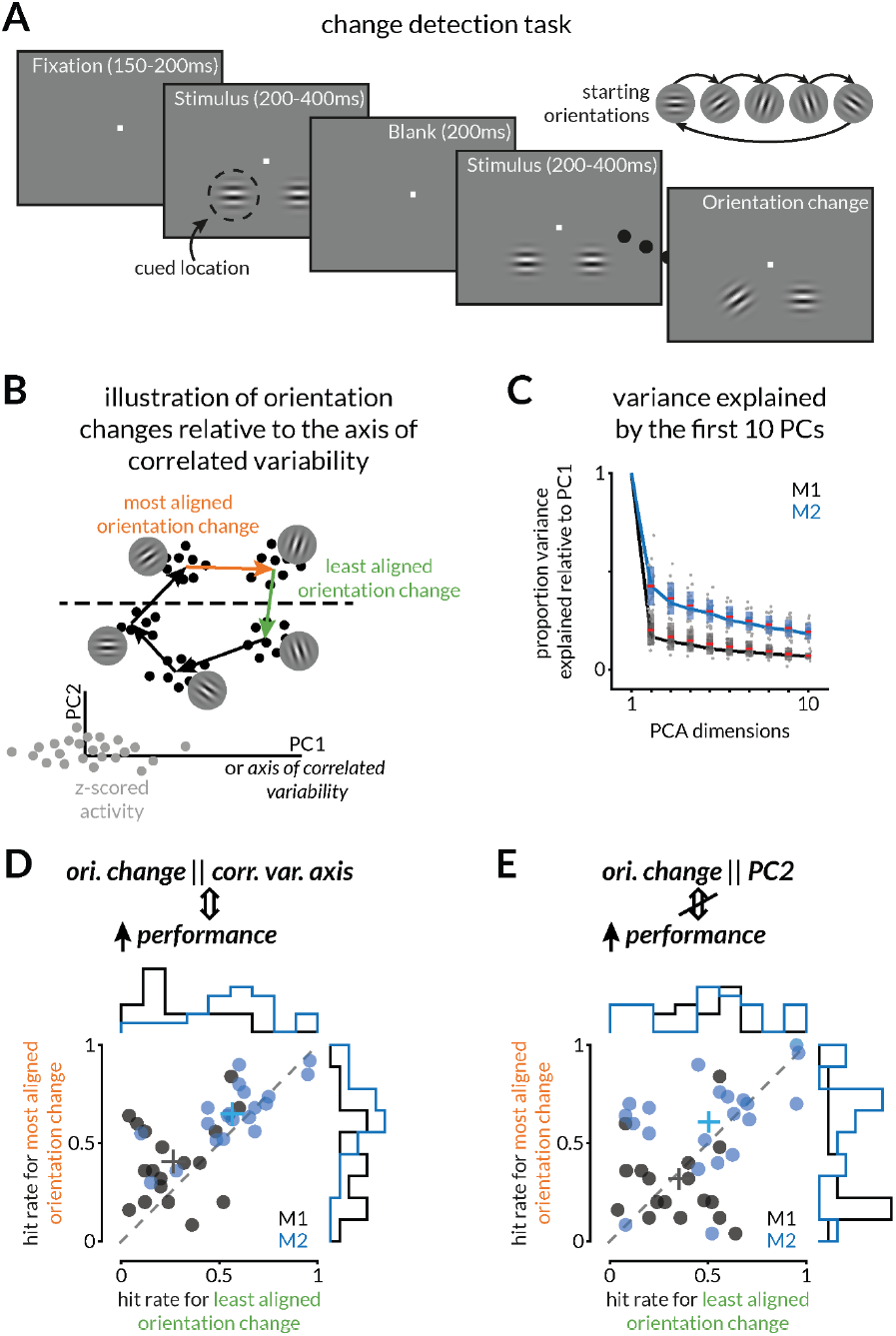
Behavioral performance in an orientation change detection task is related to the alignment between V4 orientation representations and the axis of correlated variability. A. Schematic of the orientation change detection task. Two Gabor stimuli with the same initial orientation are flashed repeatedly, and one changes orientation at a random time picked from an exponential distribution. Monkeys are rewarded for making a saccade to the location of the orientation change within 500 ms(9, 35). On each trial, the initial orientation was randomly selected between 0° and 180°, while the magnitude of the change was constant across trials. The stimulus most likely to change was cued in blocks (80% valid). We analyzed trials in which the change occurred at the cued location when that location was within the joint receptive fields of the recorded V4 neurons.B. We defined the *<u>axis of correlated variability</u>* as the first principal component of V4 activity z-scored to remove stimulus dependent variability (gray dots). Black dots represent evoked responses to stimuli with different orientations. Our hypothesis predicts that performance is better when the orientation change aligns with the axis of correlated variability (most aligned change; orange arrow) than when it does not (least aligned orientation change; green arrow).C. Variance explained by the first 10 principal components relative to the first component. The variance explained by the first component was 41.6% and 19.4% of the total variance for the two monkeys respectively.D. Validation of the prediction in B. Each point (black=monkey 1; blue=monkey 2) represents the comparison of the average hit rates on the most and least aligned orientation change conditions for one session (n=18 for monkey 1 and n=20 for monkey 2; approx. 345 and 426 trials per session; approx. 70 and 18 trials per orientation change at the attended location). Across sessions, monkeys performed better on the best aligned than on the least aligned orientation change (p=0.019, n=18 for monkey 1 and p=0.021, n=20 for monkey 2; Wilcoxon signed rank test). Marginal histograms show the distribution of hit rates across sessions. The plus signs denote mean values.E. Same as D, for the second principal component. We see no relationship between the hit rate for the orientation change most and least aligned with this dimension.

According to our hypothesis, behavioral performance should be better for orientation changes that are most aligned with the axis of correlated variability (Figure 1B, orange arrow) compared to orientation changes that are least aligned (Figure 1B, green arrow). We first verified that V4 responses to repeated presentations of the same stimulus were low dimensional (Figure 1C). We quantified alignment as the correlation of the orientation change values with the predictions of a leave-one-out cross-validated classifier trained on the neural responses projected onto the axis of correlated variability to classify the starting and change orientations. The mean classification accuracies were 0.5±0.029 and 0.641±0.01 for the least and most aligned orientation changes for monkey 1 and 0.73±0.02 and 0.96±0.01 for monkey 2 across 18 and 20 sessions respectively. Consistent with our prediction, monkeys were significantly better at detecting the orientation change most aligned with the axis of correlated variability compared to the least aligned orientation change (Figure 1D). This was not the case for the orientations sorted by alignment with the second principal component of V4 baseline activity (Figure 1E). Together, these findings show that detection performance covaries with alignment to the axis of correlated variability and that this relationship is not strongly captured by an alternative axis derived from baseline activity. This result in a well-studied task prompted us to explore our hypothesis further using a theoretical model and four additional datasets, which encompassed a wide range of stimuli and behavioral demands.

### A circuit model demonstrates that stimulus information can be read out optimally when aligned with the axis of correlated variability

To explore the neuronal mechanics required to align stimulus representations with the correlated variability axis, we constructed a network model composed of excitatory and inhibitory units with defined feedforward (*W*_*F*_), recurrent (*W*_*R*_), and linear read-out (*W*_*O*_) synaptic weights (Figure 2A). In our model, each neuron receives an external source of private (independent) trial-to-trial fluctuations, so that population co-variability only emerges through recurrent interactions between neurons (as defined by *W*_*R*_).

**Figure 2.**
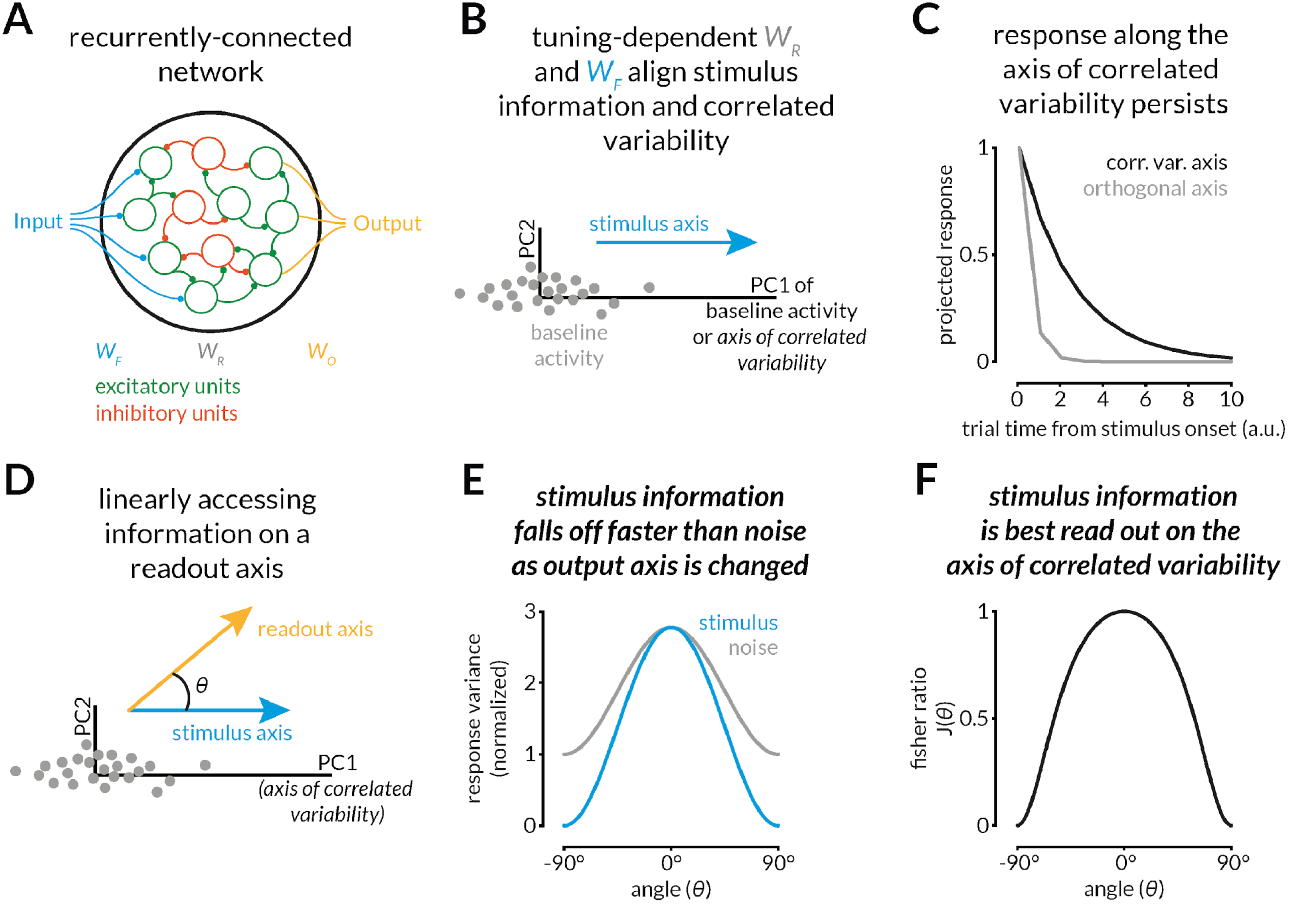
Alignment of stimulus information with the axis of correlated variability makes it optimal for readout. A. Recurrently coupled network composed of excitatory and inhibitory neuron models with rank-one feed-forward weights *WF* (blue), recurrent weights *W*_*R*_ (grey), and linear readout weights *W*_*O*_ (orange). A stimulus enters through *W*_*F*_. Independent private noise is injected into each neuron and is shaped only by recurrent weights (*W*_*R*_). The network connectivity follows Dale’s law.B. In our network, the recurrent connectivity (*W*_*R*_) is aligned with the feedforward input (*W*_*F*_), causing the stimulus axis (blue arrow) to align with the first principal component of baseline activity (PC1, grey dots)C. Once this alignment is established, fluctuations along PC1 (axis of correlated variability) decay much more slowly than along any orthogonal mode. Power on the PC1 axis (black curve) persists, whereas power on, for example, PC2 (light grey) vanishes rapidly.D. We examine a linear readout whose axis (orange) forms an angle θ with the stimulus axis (blue).E. Normalized information about the stimulus (blue) and noise (gray) as a function of *θ*. As the read-out is rotated away from the axis of correlated variability, stimulus information decreases more steeply than noise.F. Fisher discriminability 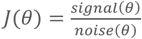 peaks at *θ = 0°*, demonstrating that, when the stimulus and noise axes are aligned, the optimal linear decoder is along the axis of correlated variability.

A wealth of data demonstrates that pairwise signal correlations are related to the magnitude of trial-to-trial pairwise noise correlations; neurons with similar tuning have stronger positive noise correlations(1, 37, 38). A likely contributor to this relationship is that recurrent coupling (*W*_*R*_) is stimulus feature dependent (*W*_*F*_), as reported in mouse V1(39, 40). Therefore, we assume that the recurrent connectivity reflects neuronal tuning and build *W*_*R*_ directly from *W*_*F*_ (see Methods). We constrained the recurrent connectivity to obey Dale’s law, the biological principle that individual neurons are either excitatory or inhibitory and therefore project connections of only one sign(41). This constraint imposes structure on recurrent interactions and shapes the geometry of population activity. Additionally, across our datasets, the axes of correlated variability computed from the baseline period and from repeated presentations of the same stimulus are well aligned (Figure S1). This independence of correlated variability from the stimulus suggests that it is not strongly shaped by feedforward inputs. We therefore model correlated variability as noise that is shaped only by *W*_*R*_, but not by *W*_*F*_.

The relationship between *W*_*R*_ and *W*_*F*_ creates a biologically realistic alignment between stimulus-evoked activity and the axis of correlated variability (Figure 2B). If the recurrent weights are unrelated to stimulus feature tuning, i.e., arbitrary and rank-one, this alignment is not guaranteed (see derivation in Supplementary Information Appendix A). Instead, this alignment comes from the relationship between *W*_*R*_ and *W*_*F*_ in our model. We also observe that after stimulus onset, response fluctuations along the axis of correlated variability persist for a longer time (Figure 2C, black curve) compared to fluctuations along orthogonal axes (Figure 2C, light gray curve). This validates previous results showing that assigning the task-relevant input to the slowest decaying dimension enables the circuit to integrate information, providing a normative rationale for the recurrent tuning(10, 42, 43). From a static encoding perspective, if a one-dimensional decoder is free to choose any direction, aligning the stimulus with the noise axis leaves the Fisher information unchanged; if the decoder is locked to a particular network mode, alignment increases information (Supplementary Information Appendix C). When the circuit must also integrate the stimulus over time, alignment becomes beneficial for any one-dimensional decoder: the slowest mode integrates the stimulus most effectively, so concentrating all stimulus power there maximizes information despite the added noise (Supplementary Information Appendix D).

We next explored how the alignment between stimulus and readout axes impacts decoding. To measure this in our model, we assumed that downstream areas could access only a single dimension of population activity (the readout axis) and that the corresponding readout weights (*W*_*O*_) remain fixed throughout the task. This is consistent with evidence that cortical circuits exploit privileged axes embedded in hard-wired projection(44). We changed the angle *θ* between the linear readout axis (Figure 2D, orange arrow) and the stimulus representation (blue arrow) and measured how much signal and noise information is aligned with the readout axis. As this angle increases, the normalized signal variance falls more rapidly than the noise variance (Figure 2E). Consequently, Fisher discriminability, a measure of decoding performance, peaks when the readout and noise axes are aligned (Figure 2F; see proof in Supplementary Information Appendix B). These results demonstrate that when stimulus information is aligned with the axis of correlated variability (a biologically relevant scenario, as correlated variability is related to signal co-tuning in essentially every study that measures both(1, 10, 45–51)), the optimal readout strategy is along the axis of correlated variability.

Our model predicts that aligning behavior with the axis of correlated variability is optimal when the relevant signal is aligned with noise. To achieve optimal coding, the core assumption of our model is that recurrent synaptic interactions (that determine the coverability axis) align with the feedforward synaptic weights (that carry signal information). This is a general assumption, consistent with the pairwise signal and noise correlations themselves being correlated, as widely reported in the literature(2, 49, 52).

### The visual feature that guides behavior is more aligned with the axis of correlated variability

We next tested the prediction that, in a situation where monkeys had to report their estimate of the value of a visual feature while ignoring other irrelevant features, the representation of the relevant feature will be aligned with the axis of correlated variability. We trained monkeys to perform a continuous curvature estimation task (Figure 3A), in which they reported the curvature of any random 3D shape while ignoring other irrelevant features, such as color, orientation, and thickness profile(53). After the stimulus was presented, the monkeys made a saccade to a location on a target arc corresponding to their curvature estimate (0 for straight, 1 for maximally curved). They were rewarded in inverse proportion to their estimation error. Both monkeys exhibited consistent curvature estimation (Figure 3B, green and blue lines; distribution of behavioral performance across sessions in Figure S3A). Their performance comparable to that of humans performing an online version of the same task (Figure 3B, pink line). The stimuli were presented within the joint receptive fields of recorded V4 neurons, which are selective for curvature (Figure 3C-D). The population representation of curvature was dependent on irrelevant shape properties (Figure 3E-F)(53), meaning that the best axis for judging the curvature of any one stimulus was different from the best for generalizing across stimuli.

**Figure 3.**
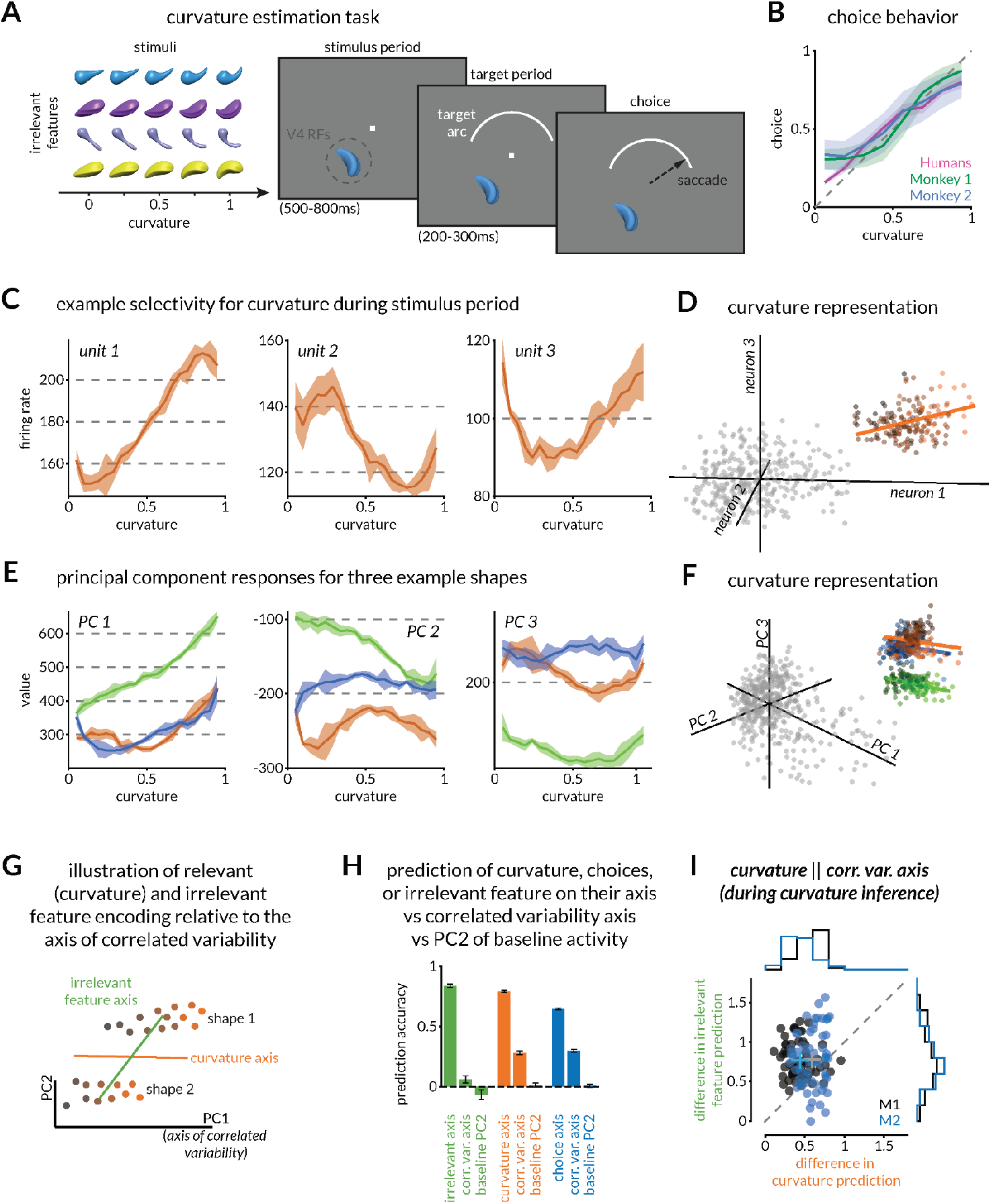
Curvature representation in V4 is aligned with the axis of correlated variability during a curvature estimation task. A. Schematic of the continuous curvature estimation task. Monkeys report the medial axis curvature of a 3D shape while ignoring irrelevant features like color, orientation, or thickness profile. Stimuli are presented within the joint receptive fields of the recorded V4 neurons. Monkeys are rewarded for making a saccade to the appropriate location on the target arc (left end for straight stimuli, right end for maximally curved stimuli). Monkeys are rewarded in inverse proportion to their estimation error.B. Behavioral responses of two monkeys (green and blue) plotted as mean reported curvature (normalized from 0 to 1) across various shapes, colors, and orientations. The shaded portion shows the standard deviation. Human participants who indicated the curvature of the same shape stimuli using a slider (pink) performed comparably to the monkeys (r=0.97 between human and monkey choices for matched curvatures). See also distributions of curvature estimation accuracies in Figure S3A.C. Example curvature tuning for three V4 multi-units for one shape. Shaded regions indicate standard error of the mean (SEM).D. Stimulus responses of the units in C plotted relative to each other (black to orange dots are curvature responses to low to high curvatures) and to their respective baseline responses (gray dots).E. Curvature tuning of the first three principal components across V4 population responses for three shapes (orange, green, and blue). The shaded region represents the standard error of the mean of the eigenvalues for each component.F. Projections onto the three PCs in E are plotted relative to each other.G. Our hypothesis predicts that the shape-general curvature representation will be better aligned with the axis of correlated variability because curvature, not other irrelevant features, drives behavior.H. Validation of the prediction in G. The curvature decoding axis is aligned with the axis of correlated variability. In the full response space, mean decoding accuracies were 0.816 and 0.862 for irrelevant feature decoding, 0.824 and 0.757 for relevant feature decoding, and 0.628 and 0.661 for choice decoding for the two monkeys respectively. When projected onto the axis of correlated variability, the mean decoding accuracies were 0.039 and 0.085 for irrelevant feature decoding, 0.238 and 0.326 for relevant feature decoding, and 0.275 and 0.325 for choice decoding for the two monkeys respectively. When projected onto the second principal component, the mean decoding accuracies were-0.137 and 0.008 for irrelevant feature decoding, 0 and 0.027 for relevant feature decoding, and-0.009 and 0.026 for choice decoding for the two monkeys respectively. Curvature decoding on the corr. var. axis was significantly higher than irrelevant feature decoding (p=6.96×10-7 and p=0.015; n=63 and n=61; Wilcoxon signed rank test).I. Per session quantification of the result in H. Each dot represents the difference between decoding performance in the full space and from responses projected onto the correlated variability axis, for the relevant (x-axis) and irrelevant feature (y-axis). Curvature (relevant feature) was significantly better aligned with the axis of correlated variability (p=2.8 × 10-4 and p=3.5×10-9; n=63 and n=61; Wilcoxon signed rank test). Conventions as in Figure 1D.

Our central hypothesis predicts that since choices depend on one feature, the representation of that feature should be aligned with the axis of correlated variability (Figure 3G). To test this, we linearly decoded either curvature or the irrelevant feature which could be color, orientation, overall shape, or some combination from the V4 population response (Figure 3H). We compared performance using the full V4 response space with performance decoding the same responses projected on to the axis of correlated variability. As predicted by our hypothesis, we could decode the relevant feature better than the irrelevant feature from responses projected onto the axis of correlated variability. This difference was not simply because relevant information is better encoded in V4: our ability to decode the irrelevant feature was slightly better than curvature in the full response space (Figure 3H). We also compared the decoding accuracy of the monkey’s choice in the full V4 response space vs when responses were projected onto the axis of correlated variability. While choice decoding was worse than curvature decoding in the full space, it was not comparably worse on the axis of correlated variability. This suggests that an axis with significant behavioral predictivity projects strongly onto the axis of correlated variability. Finally, we calculated the decoding accuracies of the irrelevant feature, curvature, and choices on the second principal component of baseline activity which on average accounted for 9.5% and 5.8% of the overall variance explained for the two monkeys respectively. We could not decode any features on this component.

To compare the alignment of the relevant and irrelevant feature representations with the correlated variability axis, we plotted the difference of decoding accuracies (from responses in the full space – projected onto the correlated variability axis) for relevant and irrelevant features (Figure 3I). For most individual sessions and for both monkeys, the decrease in decoding performance from projecting responses onto the correlated variability axis was less for relevant than irrelevant features, suggesting that the correlated variability axis was better aligned with the curvature representation than the representations of irrelevant features. We also found that curvature estimation performance was consistently better for the shape whose curvature representation was better aligned with the axis of correlated variability for monkey 2 (Figure S3B-D).

### The axis of correlated variability aligns with planning-related signals in V4 during flexible sensorimotor mapping

A corollary of our central hypothesis that correlated variability reflects task relevant information is that when the monkey maps the same visual inference to different motor outputs, the axis of correlated variability reflects the visuo-motor mapping. We consider this hypothesis because many previous studies have identified pre-motor or, more generally, behavioral planning-related signals even in primarily sensory areas, such as V4(54–60). These signals could be related to motor efference, drawing attention to context-related signals, or surround modulation and normalization relevant for the task(53). We designed a variant of our curvature estimation task in which we presented curvature variants of the same stimulus to limit the visual stimulus related variability to the curvature axis. Instead, we varied the length and angular position of the target arc so that the same stimulus curvature could be associated with different eye movements (Figure 4A). This manipulation required the monkeys to flexibly remap curvature estimates to several different saccade directions (Figure 4B).

**Figure 4.**
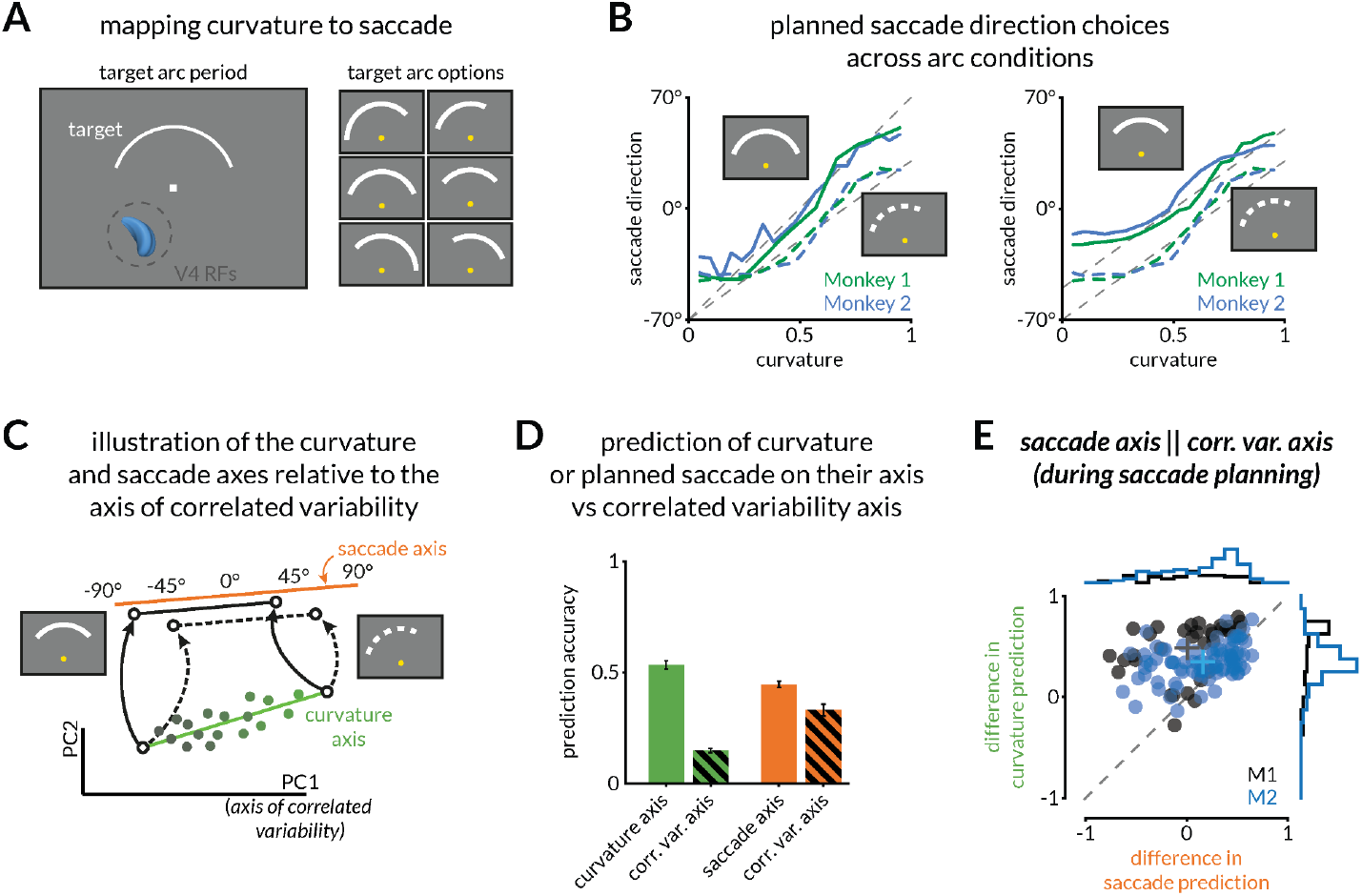
The axis of correlated variability aligns with planning-related signals in V4. A. In a subset of sessions, the length and angular position of the target arc varied across trials(53).B. Monkeys reconfigure their mapping between curvature judgement and saccade. Mean saccade directions across three arc conditions (the dashed condition is the same across the two panels for comparison) for the two monkeys (colors). The ideal judgement is indicated with the gray dashed line. Left: across two conditions with shared mapping for lower curvatures (−70°) but different mapping for higher curvatures (+70° vs +30°), saccades diverge for high curvatures. Right: across two conditions with the same arc length (100°) but different angular positions (midpoint at 0° vs-20°), psychometric curves for the two conditions have a vertical offset (32 shapes for monkey 1, 34 shapes for monkey 2).C. Our hypothesis predicts that if V4 responses are modulated such that they reflect the planned saccade, then the saccade decoding axis should be better aligned with the axis of correlated variability than the curvature decoding axis.D. Validation of the prediction in C. The saccade decoding axis is better aligned with the axis of correlated variability than the curvature axis. While both curvature (green) and planned saccade (orange) can be readily decoded from V4 responses (mean curvature prediction accuracies of 0.64±0.04 and 0.49±0.02 and mean saccade prediction accuracies of 0.38±0.03 and 0.47±0.015 across 34 and 83 shapes for the two monkeys respectively), those projected onto the axis of correlated variability are more predictive of saccade (orange dashed) than curvature (green dashed) (mean curvature prediction accuracies of 0.15±0.02 and 0.14±0.01 and mean saccade prediction accuracies of 0.38±0.05 and 0.31±0.03 across 34 and 83 shapes for the two monkeys respectively).E. The difference in curvature prediction accuracy between the curvature axis and the axis of correlated variability (y-axis) is much larger than the difference in the saccade prediction accuracy between the saccade axis and the axis of correlated variability (x-axis) (p=8.87×10-7 and p=9.98×10-6; n=34 and n=83; Wilcoxon signed-rank test). Conventions as in Figure 1D.

We previously showed that V4 population responses reformat during the motor planning period (after arc onset), such that both stimulus curvature and the planned saccade are encoded in V4(53). Because the planned saccade direction is what most directly guides the upcoming action, we hypothesized that the axis of correlated variability would align more strongly with the saccade than the curvature representation (Figure 4C). Indeed, both curvature and saccade direction could be decoded from V4 population activity. However, consistent with our hypothesis, when those responses were projected onto the axis of correlated variability, decoding performance for curvature fell more sharply than for saccade prediction (Figure 4D), suggesting that the correlated variability axis was better aligned with the saccade-related than the curvature axis.

This finding is further emphasized by comparing responses to different shapes (Figure 4E). We compared the prediction of our decoder using either the curvature or saccade direction axes with projections onto the correlated variability axis. The difference in curvature prediction accuracy between the dedicated curvature axis and the axis of correlated variability (y-axis) was substantially larger than the difference in saccade prediction accuracy between the dedicated saccade axis and the axis of correlated variability (x-axis). This result indicates that between the signals related to the visual stimulus (curvature) and motor planning (saccade), the signal pertinent to behavior is more aligned to the axis of correlated variability. Some sessions show negative differences, which likely reflect finite-sample variability in estimating high-dimensional decoders; restricting activity to one dimension (like the correlated-variability axis) can yield more stable cross-validated estimates.

### Alignment of feature representations with the axis of correlated variability improves after task learning

In the previous experiment, the monkey needed to ignore irrelevant visual features and base decisions on a single, task-relevant feature. Since V4 neurons are selective for multiple visual features, we next investigated whether we can track the alignment of a feature with the axis of correlated variability as the feature becomes relevant to choices. To test this, we trained monkeys to perform a two-alternative forced choice task that required decisions based on curvature or color on randomly interleaved trials (Figure 5A-B). On trials where the two stimuli had the same color, monkeys were rewarded for choosing the more circular (less curved) shape. When the two stimuli had the same shape, they were rewarded for making a saccade to the bluer stimulus. Both monkeys successfully made curvature- and color-based choices (Figure 5C).

**Figure 5.**
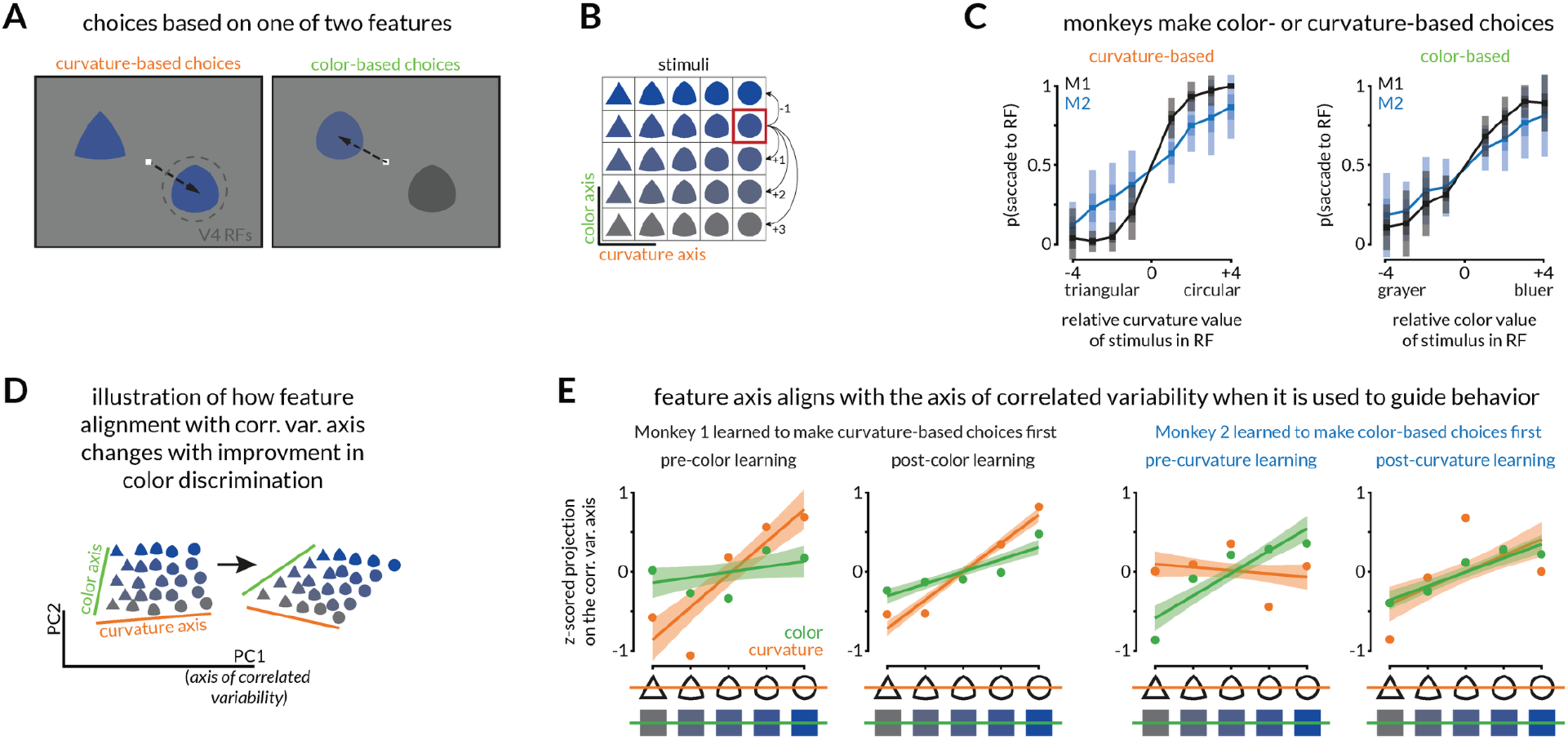
Alignment of feature representations with the axis of correlated variability improves after task learning. A. Schematic of a two-alternative forced choice task in which monkeys make either curvature-based or color-based choices(58, 61–65). If the two stimuli have the same color, the monkeys are rewarded if they make a saccade to the more circular shape. If the two stimuli have the same shape, the monkeys are rewarded if they make a saccade to the bluer stimulus. One of the stimuli is presented in the joint receptive fields of V4 neurons.B. Pairs of stimuli are selected from either the same row (curvature trials) or the same column (color trials). The relative color values are indicated on the right for an example stimulus (red outline).C. Both monkeys reliably make choices based on color and curvature on interleaved trials (30 sessions for monkey 1, 26 sessions for monkey 2; 239 and 206 color-based trials and 238 and 204 shape-based trials on average for each monkey, respectively).D. Our hypothesis predicts that the axis of correlated variability will initially be aligned only with the representation of the feature the monkeys learned to discriminate first, and then to a combination of the two feature axes that guide behavior after learning.E. Validation of the prediction in D. Early in learning for both monkeys, the representation of the feature they learned first (curvature for monkey 1 and color for monkey 2) projects on the axis of correlated variability such that it can be linearly decoded. We quantified this by testing the significance of the slope of the line relating projection onto the correlated variability axis and the stimulus value, which much be different than 0 to allow decoding (Monkey 1, 11 sessions: curvature slope p=1.01×10-10, f=62.14 vs constant model, color slope p=0.08, f=3.3 vs constant model. Monkey 2, 22 sessions: curvature slope p=0.22, f=1.54 vs constant model, color slope p=3.28×10-14, f=76.7 vs constant model.) After learning the second feature, both features project on the axis of correlated variability. (Monkey 1, 30 sessions: curvature slope p=1.19×10-51, f=376.54 vs constant model, color slope p=1.63×10-14, f=66.67 vs constant model. Monkey 2, 26 sessions: curvature slope p=1.87×10-5, f=20.49 vs constant model, color slope p=1.09×10-10, f=53.64 vs constant model)

During training, we taught the monkeys to discriminate one feature at a time. Monkey 1 learned the curvature discrimination first, and monkey 2 learned the color discrimination first. Therefore, early in training, only one of the features could guide choices. After learning, because the relevant feature changed on randomly interleaved trials, the monkeys would have to use a combined strategy (e.g. ‘choose the bluer and/or rounder stimulus’), even though one feature was uninformative on any given trial.

Our central hypothesis predicts that the neural representation of these stimuli would transform across learning to reflect a strategy that changes from discriminating one feature to considering both. The prediction is that the feature the monkey learned second, which changed from irrelevant to relevant over the course of learning, would become more aligned with the axis of correlated variability (Figure 5D). To test this, we projected V4 responses to each stimulus onto a common axis of correlated variability, computed from the baseline period before the monkeys knew whether the trial would involve curvature or color discrimination. We found that in the early sessions, only the feature the monkey learned first projected strongly onto the axis of correlated variability. After learning, both features projected strongly onto the axis of correlated variability (positive slopes in Figure 5E).

### Causal evidence: Behavioral effects of microstimulation are strongest when aligned with the axis of correlated variability

Finally, we causally tested our central hypothesis using electrical microstimulation in the middle temporal area (MT). We chose MT because the effects of microstimulation on the motion judgments have been well-established(66). As in our curvature estimation study, we trained monkeys to perform a continuous estimation task. In this case (Figure 6A), they were rewarded for correctly estimating the motion direction of a random dot kinematogram presented within the joint receptive fields of MT neurons recorded on a multielectrode linear probe. On a randomly selected subset of trials, we paired the visual stimulus with microstimulation on one of two electrodes (*long-stim*). The stimulation electrodes were chosen based on the different direction selectivity of the recorded neurons, determined in separate mapping experiments. We quantified the behavioral effect of stimulation as the change in the slope of the psychometric function relating the chosen direction to the motion direction between *long-stim* and *no-stim* conditions (Figure 6B and Figure S4A).

**Figure 6.**
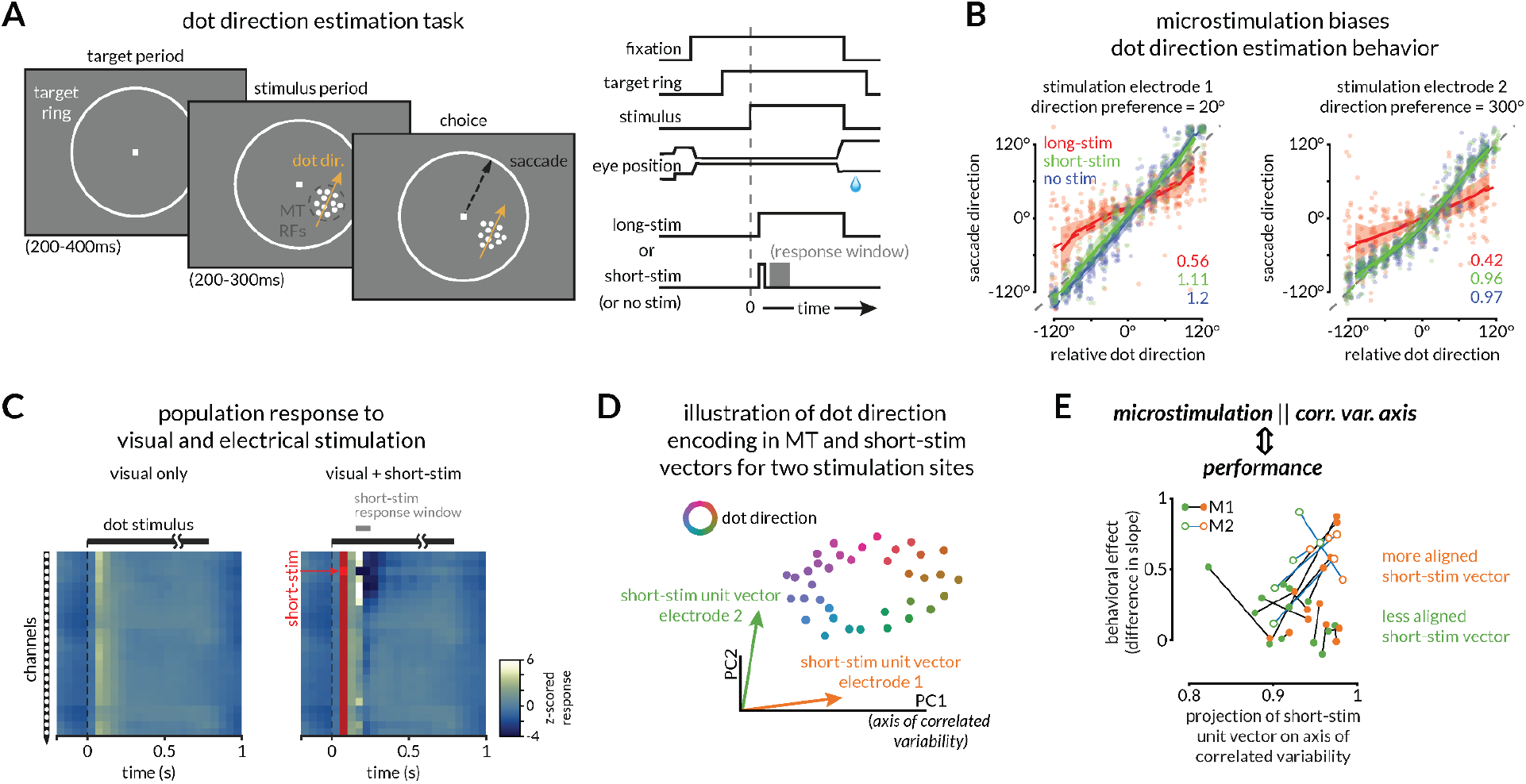
Causal test of our central hypothesis: the behavioral impact of electrical microstimulation is largest when it is aligned with the axis of correlated variability. A. Schematic of our motion direction estimation task in which monkeys estimate the direction of a random dot kinematogram displayed in the joint receptive fields of MT neurons recorded using linear probes(2, 8, 9, 19, 36, 68–70). The trial structure is illustrated on the right. After assessing the direction selectivity of recording sites in an independent mapping experiment, we chose two electrodes for electrical microstimulation. After the monkey fixated a central spot and the target ring appeared, the visual stimulus was presented. Monkeys were rewarded for accurately estimating the direction of motion by making a saccade to a corresponding point on the target ring. On some randomly interleaved trials, either a long period of microstimulation (long-stim; for the full duration of the visual stimulus) or a short microstimulation train (50 ms) was applied on one of two selected electrodes. We analyze the impact of microstimulation on MT responses during the marked analysis period (gray shaded region; 50 to 150 ms after the last stimulation pulse). We compared the monkey’s behavior on no-stim and long-stim trials. We selected the short-stim parameters to minimize the behavioral impact of stimulation while maximizing the neural impact.B. Example direction estimation behavior in no-stim, short-stim, and long-stim trials for one session. The two stimulation electrodes had preferred directions of 20° and 300°, respectively, and log-stim, but not short-stim, biased the monkey’s choices toward those directions. Here, the saccade direction for each trial is plotted against the dot direction relative to the preferred dot direction of the stimulated site. We quantified the behavioral effect as the difference in the slope of the linear fits in the no-stim and long-stim conditions (slopes are indicated in the labels at the bottom right) to the linear range (±120°) of the psychometric curves shown here.C. Example z-scored response histogram across all dot stimuli for the recording sites along the probe for no stimulation (left) and short-stim trials (right). The short 50 ms stimulation pulse train was delivered on the third site (indicated by the red box). We evaluated the effect of the stimulation by using a 100 ms response window after the end of stimulation to calculate a response vector for each of the two stimulation sites in every experimental session.D. Our hypothesis predicts that choices will be most affected by electrical microstimulation when stimulation moves neural activity along the axis of correlated variability.E. Validation of the prediction in D. For each session (two connected dots), we identified the stimulation electrode that moved MT population activity in a direction that was more aligned with the axis of correlated variability. For each electrode, we calculated the vector of population activity defined by the firing rate of each neuron on short-stim trials. We then calculated the projection of this vector onto the axis of correlated variability (x-axis; orange dots represent the electrode with the larger projection, so, by definition, orange dots are to the right of their respective green dots representing the other stimulation electrode). The impact of microstimulation on choices (difference in slope between long-stim and no-stim trials) is larger for better-aligned stimulation vectors (a majority of orange dots are above their respective green dots). See Figure S4B-D for further quantification.

We reasoned that a differential behavioral impact of stimulation across two stimulation sites must underlie a difference in how the neural effect of stimulation on those sites aligns with the axis of correlated variability. Following the logic of Moore and Armstrong, 2003(67), we designed a novel technique to measure the neural impact of microstimulation on population responses. In some stimulation trials, we truncated the microstimulation after 50 ms (*short-stim*) and analyzed responses immediately after stimulation (Figure 6C). We chose the stimulation duration for *short-stim* trials such that there was no detectable impact on behavior (Figure 6B and S4A). We calculated the alignment as the dot product between the axis of correlated variability and the microstimulation-evoked neural population response change for each electrode on short-stim trials (short-stim unit vectors in Figure 6D).

If choice behavior is preferentially influenced by activity along the axis of correlated variability, then the stimulation site that evokes responses more aligned with this axis should have a stronger impact on the monkeys’ choices (Figure 6D). As predicted, the stimulation site that pushed neuronal responses in a direction more aligned with the axis of correlated variability (orange dots in Figure 6E) also had a larger impact on behavior on long-stim trials. In other words, the slopes of the lines connecting the metrics for the two electrodes are largely positive. We quantified the relative projections of the impact of microstimulation from the two sites in each session onto the correlated variability axis by calculating the slopes of the connected lines in Figure 6E (Figure S4B). The distribution of actual slopes was significantly skewed compared to the distributions of analogous slopes reflecting projections on the axis representing either the mean stimulus-evoked firing rate or the dominant stimulus-evoked activity axis (compare Figure S4B, S4C, and S4D). This provides causal evidence that neural response fluctuations along the axis of correlated variability, but not along alternate axes derived from stimulus-evoked activity, are strongly related to behavior.

Together, our results show that the axis of correlated variability does not merely reflect noise to be ignored but instead aligns with—and may amplify (or be a byproduct of response amplification)—the sensory and decision-related signals that guide behavior. This framework reconciles the tight relationship between correlated variability and behavior with the theoretical possibility of decoding information in its presence, pointing to a mechanism by which perception and action can remain flexible, robust, and efficient.

## Discussion

Our results demonstrate that shared, trial-by-trial response fluctuations in the visual cortex are not simply ignored by downstream decoders. Instead, shared variability reflects the information that guides behavior and tracks changes in task demands. Using complementary correlative, theoretical, and causal approaches and several visually guided tasks, we demonstrated that:

1. Change detection is most accurate when stimulus representations vary along the axis of correlated variability;
2. When feedforward coupling (signal) and recurrent coupling (noise) are aligned in the same circuit, decoding along the axis of correlated variability is optimal;
3. The axis of correlated variability flexibly reflects task demands, from sensory features to action plans;
4. When perception and action are dissociated, the axis of correlated variability favors motor intent over sensory evidence;
5. Causal manipulations are most effective when aligned with the axis or correlated variability.

Together, these findings support our central hypothesis: the axis of correlated variability shares a substantial subspace with the neuronal population activity that is read out to guide behavior. These findings bridge a long-standing gap between theoretical accounts of information-limiting correlated variability and empirical observations linking response variability to attention, learning, and behavior. They suggest that neuronal variability is not a nuisance, simply tolerated by the brain. Rather, it reflects the substrate of perception and decision-making.

### Why measure correlated variability?

In recent years, a staggering number of studies have linked various aspects of flexible, sensory-guided behavior to a remarkably simple measure of shared variability: the mean correlation between the spike count responses of pairs of neurons to repeated presentations of the same stimulus. Such correlated variability is modulated by nearly every process that affects perception, including attention, adaptation, learning, task switching, arousal, and stimulus contrast(20, 48), and is linked to behavior on a trial-by-trial basis(1, 10, 16, 17, 46, 48–51). The prevalence of this relationship suggests a deeper role: that correlated variability provides a window into the neural computations that support cognition.

Specifically, three lines of prior evidence guided our thinking. First, signal and noise correlations are inextricably intertwined. Neurons with similar tuning tend to have higher noise correlations(10, 42, 43), likely due to an underlying shared circuit structure(54, 55, 68, 71–74). Second, we previously showed that in a change detection task, the axis of correlated variability reflects all the choice-predictive information in V4, even when attention reduces mean correlations. Third, the flexibility of this relationship across stimuli, tasks, and behaviors mirrors the flexible contribution of the visual cortex to visually guided behavior. The responses of neurons in areas like V4 and MT, and their correlated variability, are modulated by attention, task switching, reward, and motor planning(16, 23, 26, 35, 68, 75), and yet a stable linear decoder can often predict behavior across those contexts(16, 26, 28). We show that the axis of correlated variability aligns with stimulus feature representations (like curvature axis in Figure 3 or color axis in Figure 5), or dimensions associated with other task-relevant information (like pre-motor signals in Figure 4 or the influence of electrical microstimulation in Figure 6) that are related to behavior. These results suggest representations of task-relevant adaptively align with the axis of correlated variability.

### Reconciling the links between noise, information coding, and performance on perceptual tasks

Correlated variability is reliably associated with performance on demanding perceptual tasks. Across species, brain areas, and tasks, decreases in correlated variability have been observed alongside good performance. This relationship is typically interpreted as reflecting information encoding: reductions in correlated variability are presumed to reflect less population-wide noise, which improves readout(16, 26, 28). However, both theoretical and empirical work challenge this interpretation. Theoretical work has demonstrated that optimal decoders can effectively ignore low-dimensional shared variability without compromising performance(14, 33, 35), questioning the need to suppress correlated variability in the first place. Consistent with this theory, we and others have empirically shown that attentional reductions in noise correlations do not improve population-level stimulus encoding(26, 27).

This raises a paradox: if shared variability does not limit information, why does it change with cognition, and why is it so closely linked to behavior?

Rather than viewing correlated variability primarily as noise to be suppressed, our results suggest a different interpretation. We propose that correlated variability reflects the content of the population representation that is most relevant for guiding behavior. In this framework, the same circuit interactions that shape how behaviorally relevant signals are amplified also shape the geometry of shared variability. As a result, correlated variability covaries with performance not because it limits information, but because it reflects the population dimensions that are most strongly involved in guiding behavior.

We assess population activity using a geometric framework in which stimulus- and task-related signals are represented in low-dimensional subspaces of a space where each dimension represents the activity of one neuron in the population(76). Across several datasets, we found that the dominant axis of correlated variability aligns with the population representation of the variable that is most relevant for behavior in that task context. Importantly, our results do not implicate a particular kind of signal (like pre-motor or preparatory signals) that must align with the axis of correlated variability, nor do they prescribe a brain area in which this alignment is most potent for behavior. Rather, it suggests that when a population represents a combination of variables that guides decisions, the largest mode of shared variability reflects that representation.

Our findings also situate this framework within a broader body of empirical and theoretical work on structured population activity. They are consistent with prior studies linking structured correlated variability to flexible sensorimotor mappings across different areas and species(6, 15, 17, 18, 45, 77–79). Notably, studies have shown that the structure of correlated variability approximates the underlying stimulus tuning distributions(80) and is related to the statistically optimal expectations of stimulus features across development(81). These results and ours confirm theoretical perspectives proposing that spontaneous and task-related activity reflect structured representations of sensory features and expectations(10, 42, 82, 83). In Bayesian and probabilistic inference frameworks, shared variability can reflect uncertainty or beliefs about task-relevant latent variables, and learning-dependent changes in covariance structure may track shifts in those beliefs(84). Our results are compatible with this perspective: as populations of neurons across brain regions perform probabilistic inference to coordinate task-relevant information across areas, the alignment between the relevant sensory variables and the axis that tracks the largest shared variability is a signature of the inference process(82–87).

A complementary line of theoretical work addresses how such structured variability can coexist with efficient coding. Recent analyses suggest that high-dimensional population codes can contain dominant modes of shared variability alongside weaker modes without degrading information transmission(88, 89). Within this broader computational context, our findings provide empirical support for the idea that the dominant axis of shared variability is systematically related to behaviorally relevant population signals across tasks and brain areas.

### A biologically plausible implementation

Traditionally, models of cortical circuits treat signal and noise as separable quantities. In that scenario, low-dimensional noise can be easily averaged out or ignored so that it is irrelevant to behavior(10, 42, 43). This is contrary to the results presented here, suggesting that, instead, correlated variability reflects the information used to guide behavior.

Our results may simply be a consequence of a biological constraint: that signal and noise emerge from the same circuit. The same connectivity that shapes stimulus representations also determines the structure of shared variability(90). When these are related, as is widely reported empirically, the axis that accounts for most shared variability will also carry the most signal.

To formalize these ideas, we developed a circuit model that links recurrent network dynamics to the structure of shared variability (Figure 2). When the feedforward and recurrent connectivity are related, the stimulus representation aligns with the dominant recurrent mode—the axis of correlated variability (Supplementary Information Appendix A). Because this mode is also the slowest, projecting stimulus drive onto it enables the circuit to integrate evidence over time (Figure 2C), providing a normative rationale for signal–noise alignment(42). Once this alignment is established, the optimal one-dimensional readout is along this common axis, because signal power falls faster than noise variance as the readout rotates away (Figure 2E-F; Supplementary Information Appendix B). The model thus predicts that, for a given task, the stimulus, noise, and readout axes should coincide, which is consistent with our experimental finding that behavior tracks the axis of correlated variability.

### Causal manipulations for linking neural population representations to behavior in primates

The primary challenge for using causal manipulations to study the relationship between behavior and correlated variability is the need to measure how manipulations affect neuronal populations. While imaging has been used to measure how electrical stimulation affects neurons in mice(91–95), it is challenging and uncommon to make these measurements using physiology and in monkeys.

The methods we used for measuring the impact of electrical microstimulation on surrounding populations (Figure 6) are broadly applicable for at least three reasons. First, microstimulation remains a uniquely effective causal manipulation for eliciting behavioral changes, especially in primates. Second, microstimulation leverages the functional organization of the cortex: by varying simple parameters, such as current amplitude, one can easily adjust the number and variety of affected neurons. Finally, microstimulation remains essentially the only causal method for assessing the function of small groups of neurons during human neurosurgery or for prosthetics in humans^80–84^. Understanding the relationship between electrical stimulation, neuronal population activity, and behavior, therefore, has implications both for basic science and translational research.

Here, electrical microstimulation in MT (Figure 6) provided a causal test of our hypothesis. Stimulation perturbed population responses along different axes (Figure S4B-D); the size of the behavioral impact was correlated with the extent to which those perturbations aligned with the axis of correlated variability. This result supports the idea that downstream circuits are sensitive to changes along this axis.

### Correlated variability as a handle on cognition

Although correlated variability has primarily been studied from a basic science perspective, our results have translational implications. Correlated variability can be modulated by cognitive states, pharmacological agents, and direct circuit manipulations. If it indeed reflects the information flow from sensory cortex to decision-making circuits, the possibilities for measuring and changing it are endless. Indeed, it is straightforward to modulate correlated variability using existing pharmaceuticals as well as by natural cognitive processes. Our results suggest that correlated variability may provide a powerful biomarker and a potential intervention target for repairing or enhancing perception in health and disease.

In sum, our results reveal that the axis of correlated variability is not just a nuisance byproduct of cortical computation. It is a window into the aspects of neural population activity that flexibly link sensory representations to behavior. By reframing variability as an adaptive feature rather than a limitation, this work offers new insight into the neural basis of perception and points to shared variability as a promising target for understanding and influencing cognition.

## Acknowledgements

We are grateful to K. McKracken for providing technical assistance, to John Maunsell for comments on an earlier version of this manuscript, and for helpful comments and suggestions regarding data analysis. We would also like to thank Dr. Lori Holt and Christi Gomez from the Holt lab for guidance and support with the online human psychophysics experiments. This work is supported by Eric and Wendy Schmidt AI in Science Postdoctoral Fellowship (to R.S.), the Simons Foundation (Simons Collaboration on the Global Brain award 542961SPI to M.R.C. and B.D.), and the National Institutes of Health (awards K99EY035362 to R.S., R01EY022930 and RF1NS121913 to M.R.C., R01NS133598 and R01NS137194 to B.D., CRCNS-R01EY034723 to M.R.C. and B.D., and training grant R90DA060338 to Y.X. and B.D.). B.D. benefited from Physics Frontier Center for Living Systems funded by the National Science Foundation (PHY-2317138), and support from the National Institute for Theory and Mathematics in Biology (Simons Foundation award MP-TMPS-00005320 and NSF award DMS-2235451). Y.X. is supported by a NeuroData Discovery Grant from the Kavli Foundation.

## Author Contributions

Study conception: RS, BD, and MRC.

Electrophysiological data collection: Figure 1: AMN and DAR, Figure 3-5: RS, Figure 6: DAR.

Computational and statistical analysis and visualization: RS.

Rate model and related analysis in Figure 2: YX.

Original writing: RS and MRC.

Reviewing and editing: All authors.

Project administration and supervision: BD and MRC.

## Data and Code Availability

The data and code used to perform the analyses and generate the figures in this study has been deposited in a public GitHub repository https://github.com/ramanujansrinath/corrvar. Further information and requests for raw data should be directed to and will be fulfilled by Ramanujan Srinath.

## Materials and Methods

### Experimental Model Details

We analyzed data from four different datasets, detailed below:

- Change detection experiment (Figure 1): Subjects were two adult male rhesus monkeys (*Macaca mulatta*, 8 and 10 kg).
- Curvature estimation experiment (Figure 3-4): Subjects were two adult male rhesus monkeys (*Macaca mulatta*, 8 and 9 kg).
- Curvature-color 2AFC experiment (Figure 5): Subjects were two adult male rhesus monkeys (*Macaca mulatta*, 11 and 10 kg).
- Dot direction estimation experiment (Figure 6): Subjects were two adult male rhesus monkeys (*Macaca mulatta*, 8 and 10 kg).

Before training, all monkeys were surgically fitted with a customized titanium head-post (Crist Instruments Co., Hagerstown, MD). Monkeys were then trained to perform their respective tasks until satisfactory performance was reached. To enable electrophysiological recordings, we chronically implanted monkeys with 96-channel microelectrode arrays (Blackrock Neurotech, Salt Lake City, UT) for recordings in V4 or acute recording chambers over area MT. For the acute recordings, we used 24 or 32-channel linear probes (V- and S-probes; Plexon Inc., Dallas, TX) positioned using grids (Crist Instruments Company Inc., Hagerstown, MD) and advanced using a hydraulic microdrive (Kopf Instruments, Tujunga, CA). All animal procedures for experiments in Figures 1, 3, 4, and 6 were approved by the Institutional Animal Care and Use Committees of the University of Pittsburgh and Carnegie Mellon University, where the electrophysiological and psychophysical data were collected. Animal procedures for experiments in Figure 5 were approved by the Institutional Animal Care and Use Committees of the University of Chicago. Additionally, all data and analyses in this study are reported in accordance with ARRIVE animal use and reporting guidelines.

#### Curvature estimation experiment (Human version)

We conducted an online psychophysics experiment in which humans reported the curvature of 3D shapes identical to those used in the monkey version shown in Figure 3. These experiments were approved by the Institutional Review Board (IRB) of Carnegie Mellon University. Participants provided informed consent electronically before beginning the task. The consent form described the study purpose, procedures, risks, benefits, compensation, and confidentiality protections, and required participants to confirm that they were 18 or older and met the study’s eligibility criteria. Only individuals who affirmatively indicated their consent were allowed to proceed with the experiment. The experiment was conducted on the online psychophysics platform, Gorilla (www.gorilla.sc)(96), and 40 human subjects (ages 19-64; average age 28) were recruited via Prolific (www.prolific.co) between February 7 and 11, 2022. Participants were randomly assigned to one of two groups that each reported the curvatures of five random shapes and their orientation and curvature variations.

Because this study was conducted online, we included additional measures to ensure task compliance. We collected data from three questionnaires, which included compliance, demographics, and feedback, per the IRB protocol. Additionally, we detected their monitor size and scaled images accordingly. We confirmed the image size via a standardized credit card size check protocol.

### Behavioral, Electrophysiological, and Computational Methods Details

#### Common experimentation apparatus

Visual stimuli were displayed on a 24” ViewPixx monitor (1920 × 1080 pixels; 120 Hz refresh rate) or a CRT monitor (1024×768 pixels; 120 Hz refresh rate), both calibrated to linearize intensity, placed 52-60cm away from the monkey. The behavioral experiments (behavioral monitoring, visual display, reward delivery, experimental, and data syncing) were performed using custom MATLAB software and the Psychophysics Toolbox(97). A square marker on the screen was flashed at the onset of stimuli, which was captured by a photodiode to synchronize the stimulus display with data acquisition. We monitored eye position using an infrared eye tracker (EyeLink 1000 Plus; SR Research). Spiking activity, local field potentials, eye position, and task events were recorded at 30K samples/s using either CerePlex E headstage and CerePlex amplifier (Blackrock Neurotech, Salt Lake City, UT) or Trellis software and Ripple recording hardware (Ripple, Salt Lake City, UT).

#### *Common behavioral, electrophysiological recording, and analysis considerations* Filtering and spike thresholding

We band-pass filtered (250-5000 Hz) the raw electrical activity (acquired at 30K samples/s) and detected threshold crossing timestamps on each recording channel with a manually set threshold (2-3x RMS signal value for each channel). These spiking events, the original raw data, and stimulus-locked photodiode activity were all saved at 30 KHz, and the eye tracking signals were saved at 2 KHz. In this study (as in previous studies from which these data originate), we did not distinguish between sorted single-unit and multi-unit activity.

#### Baseline response

In all datasets, we calculated the baseline response during the trial epoch after stable fixation and before the onset of the visual stimulus. During this time, the monkey was fixating on a central dot displayed on a gray screen. In most experiments, we varied this duration between 150 and 250 ms, drawing randomly for each trial from a uniform distribution to prevent the monkey from learning the precise timing of the task. We calculated the spike rate for each unit during a fixed, minimum duration for each session. Stimulus response: We calculated the stimulus-evoked spike rate after a latency of 50 ms (to allow for latency of responses in V4 and MT) during the stimulus display epoch for trials with stable fixation. The details of stimulus durations were specific to each experiment and are detailed below.

#### Neuron inclusion

For all population-level analyses, we only included units if their average stimulus-evoked response was at least 10% higher than during the baseline, gray-screen period.

Trial inclusion: In all datasets, we analyzed only trials in which the monkey completed either correctly or incorrectly. We excluded trials during which the monkey made a premature saccade to break fixation or those where we detected spurious electrical noise artifacts in the neural recordings.

#### Calculation of the axis of correlated variability

We performed Principal Components Analysis (PCA) on the baseline activity for each session and defined the axis of correlated variability as the first principal component of this activity. In previous studies(2, 12, 36), we have calculated this axis, spike count correlations, or the covariance matrix using repeated presentations of the same stimulus. We repeated this analysis for all three datasets and found that the correlated variability axes calculated from baseline and evoked responses were extremely similar (Figure S1). Since the baseline response can be measured during every trial, we opted to calculate the axis of correlated variability using this response.

#### Change detection experiment

We analyzed data from a previously published dataset in which monkeys performed a cued attention change-detection task while we recorded neuronal activity from area V4(34, 35). Briefly, we analyzed data from a subset of 20 sessions during which the monkeys performed a variant of the change detection task with multiple starting orientations but a constant change amount. During instructional trials, monkeys fixated a central dot while we flashed a single Gabor at the location where the orientation was likely to occur. During subsequent trials, monkeys maintained central fixation while two peripheral Gabor patches were flashed repeatedly (200 ms on, 200-400 ms off). These Gabors had an orientation drawn from a limited set of either [0°, 45°, 90°, 135°] or [0°, 36°, 72°, 108°, 144]. At a random time, the orientation of the Gabor at the cued location changed, and the monkeys were rewarded for making a saccade to the changed stimulus. The orientation change amount was constant throughout the session at either 45° or 36°, depending on the starting orientation set. Spatial attention was manipulated in blocks of trials, each starting with a set of instructional trials. In this study, we only analyzed 80% of the trials in which the change occurred at the cued location and orientation differences for which performance exceeded 20% across all sessions. Mean hit rate for the most and least aligned orientation change conditions were 41% and 26.5% for monkey 1 and 65% and 56.5% for monkey 2. The stimulus responses were calculated for each flash during the 60-130ms epoch after stimulus onset. Average behavioral performance for each session was calculated as the percentage of hits in detection for each orientation change. Miscellaneous experimental details of session inclusion, mean firing rates, receptive field mapping, etc., can be found in the original publications(21, 34, 35). As in previous studies using this task, we calculated the axis of correlated variability using evoked responses to repeats of the same visual stimulus. This strategy is advantageous because it quantifies correlated variability using the decision formation period. Because our other tasks do not include many repeats of the same stimulus and because it is a stronger test of our hypotheses, for our other data sets, we calculated the axis of correlated variability using baseline responses from the initial period where the monkey fixates a blank screen. We compared axes calculated from stimulus and baseline responses in Figure S1 and discovered that the two are highly similar.

#### Curvature estimation experiment

We analyzed data from a previously published dataset in which monkeys performed a continuous curvature estimation task while we recorded activity from area V4(53). Briefly, we analyzed data from a subset of 82 sessions during which monkeys fixated a central dot presented on a gray screen while a randomly generated 3D stimulus was shown in the joint RF of V4 neurons for 550-800ms. For each session, 3-6 stimuli were drawn from a set of 120 base shapes that vary in overall shape (thickness profile, gloss, twist, length, out-of-plane rotation, etc.) or in-plane orientation only or color only. The curvature of the selected shapes was varied, drawn from a uniform distribution across trials in 20 (monkey 1) or 10 (monkey 2) steps. After the stimulus presentation period, a target arc was presented in the upper hemifield. In a majority of sessions, a 140° target arc was presented centrally (82 sessions). In a non-overlapping set of sessions (57 sessions; relevant for analyses in Figure 4), either the angular position (0° or ±20°) or the length of the arc (100° or 140°), or both, were randomized across trials. After the presentation of the arc, the monkeys made a saccade to the arc to indicate their curvature inference. These saccade directions were converted to curvature inference by mapping the possible saccades (−70° to 70° on the 140° centrally presented arc, say) to a scale of 0-1. The reward amount fell linearly along the arc centered on the correct curvature value up to a threshold (±0.1), after which it fell to 0. We calculated the stimulus-evoked firing rate for each unit during an epoch of 50-550 ms after stimulus onset to allow for V4 response latencies. We calculated the arc-evoked firing rate during an epoch of 0-150 ms after the onset of the arc. Average behavioral performance was calculated for each shape across curvature variations as one minus the average absolute error in curvature judgment. Details about stimulus construction, behavioral timing, reward landscape, and RF mapping can be found in the original manuscript(53).

#### Curvature-color 2AFC experiment

We analyzed behavioral and neural data from V4 in a two-feature discrimination two-alternative forced-choice (2AFC) experiment. Data from monkey 1 were analyzed and discussed as part of a previous manuscript(53). We added additional data from monkey 1 and repeated experiments in monkey 2. Briefly, 25 shape stimuli were created by varying the color between gray and blue (isoluminant) in 5 steps and the ‘curvature’ of the stimuli in 5 steps. The curvature was varied by creating homeomorphs of stimuli between an equilateral triangle and a circle using linear interpolation. After the monkey fixated a central dot for 150-250 ms, two shapes that either shared a common curvature value or a common color were sampled from the grid of 25 stimuli and presented in opposite hemifields (with one stimulus location overlapping the joint RFs of V4 neurons). The stimuli were displayed for 200-250 ms, after which the fixation point was removed, serving as a go cue for the monkey to make a saccade to one of the two stimuli. The monkey was rewarded with a drop of juice for selecting the stimulus that was bluer and more circular. We calculated the stimulus response during a window of 50-200 ms after stimulus onset for each trial. We measured behavioral performance (psychometric) curves by calculating the difference in value of the visual feature (color or curvature) that the monkey can use to guide behavior between the stimulus in the RF and the stimulus in the opposite hemifield. This difference could take eight values between-4 and 4, excluding 0, as we did not present two identical stimuli. Psychometric curves in Figure 5 depict the probability that the monkey chose the stimulus in the RF for each comparison.

#### Dot direction estimation experiment

Our central hypothesis predicts that external perturbations that are aligned with the axis of correlated variability would have a larger behavioral effect compared to perturbations that are not. We designed a new causal test: we compared the behavioral and neural impacts of electrical microstimulation on two recording sites in the middle temporal area (MT) while monkeys performed a continuous random dot direction estimation task (Figure 6).

Monkeys fixated a central dot on a gray background before a target ring was presented for 200-400 ms. Then, as the monkeys continued to fixate, a dynamic random dot kinematogram was displayed at a location that overlapped the RF of the MT neurons. Monkeys were rewarded for making a saccade to a location on the target ring that corresponded to the direction of the kinematogram, not where the target ring intersects the dot motion vector. Behavioral accuracy was calculated as the slope of the linear relationship between the actual dot direction and the monkey’s saccade direction. Unlike the other datasets in this manuscript, the neural data in this experiment were recorded using linear probes with 24 or 32 channels, rather than microelectrode arrays; however, the data were acquired, pre-processed, and analyzed in the same manner. The probes were inserted such that recorded MT units had highly overlapping RFs but different direction tuning preferences. The RFs and direction tuning preferences were measured using an independent experimental protocol to guide stimulus placement and selection of microstimulation sites. In subsets of trials, one of two pre-selected contacts was microstimulated using a biphasic, 200 Hz pulse train with an amplitude ranging between 20 and 40 µA.

Quantifying the behavioral effect of microstimulation: During *long-stim* trials, this pulse train temporally overlapped the visual stimulus. We used these trials to measure the behavioral effect of microstimulation on that channel, calculated as the difference of the slope relating the dot direction to the saccade direction between the *long-stim* and *no-stim* trials. We also quantified the behavioral effect of *short-stim* (Figure S4A).

Quantifying the neural effect of microstimulation: During *short-stim* trials, the pulse train started ∼140 ms after stimulus onset and lasted for 50 ms. We quantified the effect of this short microstimulation train by calculating the stimulation-evoked spike rate for each recording site during the epoch 50 ms after the termination of the last pulse and lasting 100 ms. (We tried 50 ms, 100 ms, and 150 ms epoch durations and found qualitatively similar results.)

Other details of RF mapping, session selection, and behavioral training and timing can be found in a manuscript describing related data(67).

#### Curvature estimation experiment (Human version)

Since the curvature estimation task is bounded on both ends, subjects routinely overestimate lower curvatures and underestimate higher curvatures. Additionally, we aimed to eliminate the possibility that idiosyncrasies in learning history contributed to any systematic variation in behavioral performance between the two monkeys. It was not feasible to repeat the curvature estimation experiment with many more monkeys, so we designed an online human psychophysics experiment to be run on a large cohort of people, providing us with a baseline to compare monkey performance to. After a set of compliance- and demographics-related questionnaires, humans performed a slider-based version of the curvature estimation experiment where a horizontal slider was presented with a stimulus image. The initial position of the slider was randomly set across trials, and humans had to use their computer mouse to select a value between 1 and 10 in steps of 0.1. The stimulus images were drawn from the same image set used for the monkeys. We divided our human cohort (n = 40) into two groups. Each group was shown 20 curved variations of 20 shapes (five base shapes at four orientations each). After each trial, the correct curvature was indicated on the slider along with their choice. The maximum time allowed per trial was 4 seconds. We recorded each choice and reaction time. No subject was excluded from analysis.

#### Network modelling

Figure 2 illustrates a linear-rate network that satisfies the rank-one and alignment assumptions, as well as respecting Dale’s law. First, we analyze a simple rank-one circuit whose feed-forward vector and recurrent Perron mode are identical, thereby aligning the noise and stimulus-response directions. Next, we embed an explicit excitatory–inhibitory structure into the same rank-one scaffold, renormalize it so the leading eigenvalue remains, and thereby obtain the Dale-compliant network that produces Figure 2.

### Dynamics and general notation

We modelled a linear-rate network of *N* = 120 units. Their activities r(*t*) ∈ ℝ^*N*^ obey

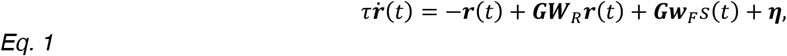

with τ = 1.

- *s*(*t*) ∈ {0,1} is a binary stimulus,
- 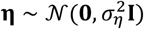 with σ = 1,
- **G** = diag(*g*_1_,…, *g*_*N*_) ≻ 0 are the neuronal gains; here we set **G** = **I**,
- **w**_F_ is the feed-forward drive,
- **W**_R_ is the recurrent weight matrix.

Whenever the spectral radius ρ(**W**_R_) < 1, the steady-state response is Gaussian (the derivation can be found in Appendices A and C of the Supplementary Information):

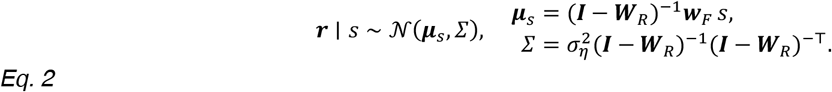

We constrained the spectral radius (i.e., the magnitude of the largest eigenvalue) of the recurrent connectivity matrix to be less than one. This condition ensures that recurrent amplification remains bounded and that the network dynamics are stable.

The signal vector Δμ = μ_1_ – μ_0_ and the linear Fisher ratio

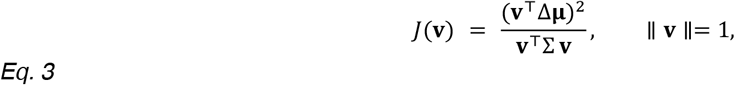

quantify information along any unit decoder **v**.

### Rank-one symmetric network

We start with the simplest rank-one construction for the feed-forward and recurrent connections. Feed-forward drive.

Let *N* = 120 be the population size; to simplify the exposition we choose a stimulus acts only on neuron 1 through the unit vector **w**: = **e**_1_ = (1,0,…,0)^⊤^. In the next section we relax this condition in our Dale’s law compliant network used to produce Fig. 2. With a stimulus-amplitude difference of β = 1.0, we set

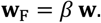

Recurrent weight matrix.

The simplest way to amplify signal and correlated noise along the same axis is to use a rank-one outer product whose Perron eigenvector coincides with **w**. Choosing a spectral radius of ρ = 0.4 gives

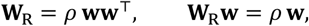

so **w** is both the stimulus and the noise direction. This construction satisfies the stimulus–noise alignment condition; Appendix A of the Supplementary Information details the general conditions under which the stimulus-response axis aligns with the dominant noise (slow-mode) axis. Because **W**_R_ is rank 1, the Sherman– Morrison identity yields (see Proposition 1 in Supplementary Methods for the proof):

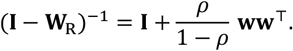

With a unit private-noise standard deviation σ_η_ = 1.0, Eq. 2becomes (see Proposition 2 in Supplementary Methods for the proof):

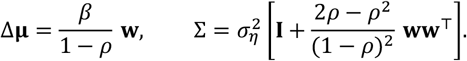

Any unit decoder can be parameterized by an angle θ as **v**(θ) = cosθ **w** + sinθ **u**_⊥_, where **u**_⊥_ ⊥ **w** is an arbitrary orthonormal complement. Substituting the expressions above into the Fisher-ratio definition Eq. 3 gives for our symmetric matrix *W*_R_ the following:

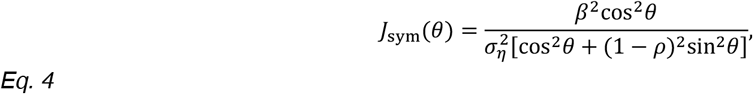

which reaches its unique maximum at θ = 0^∘^, i.e. when the decoder is aligned with the common stimulus/noise axis **w**. For general **W**_R_ and **w**_F_ the noise axis is still aligned with the stimulus-response axis, the single-axis decoder remains optimal at θ = 0^∘^ (shown in Appendix B of the Supplementary Information).

### Biologically plausible implementation: a Dale-compliant rank-one network

We now redesign the rank-one network framework so that the network respects Dale’s law and has a feedforward stimulus structure that better mimics how a tuned population is driven. This is the version that generates the results presented in Fig. 2.

Magnitude profile.

We first place the 120 neurons along a line *x*_*i*_ = −1 + 2(*i* − 1)/(*N* − 1) and assign each neuron:

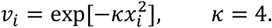

We normalize the vector as **v**: = **v**_mag_/∥ **v**_mag_ ∥ so that ∥ **v** ∥= 1. This choice makes the recurrent weights largest near the middle and smaller toward the ends, producing a smooth “bump’’ profile.

E/I sign pattern.

We divide the neurons into excitatory (80 %) and inhibitory (20 %) groups. Rows 1–96 receive a + sign (excitatory) and rows 97–120 a − sign (inhibitory). Element-wise multiplication produces the signed vector

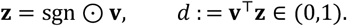

Because most entries are positive, *d* is close to 1 (*d* ≈ 0.985 with the default parameters). We will divide by *d* in the outer product below to keep the Perron eigenvalue exactly ρ, ensuring that the Dale’s law compliant network inherits the same overall gain and time-scale as the simpler symmetric network.

Recurrent matrix.

Combining magnitude and sign vectors yields a single rank-one matrix

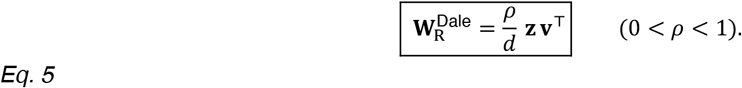

*Why the factor* 1/*d?* Right-multiplying by **z** gives 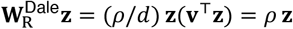. Thus **z** is the Perron eigenvector with exactly the same eigenvalue ρ used in the simple symmetric network.

Feed-forward drive.

To mimic the symmetric network we inject the stimulus *along* the Perron vector:

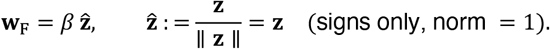

Equivalence to the symmetric rank-one model.

Although it looks different, 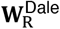 is still rank 1 and shares the same non-zero eigenvalue ρ. Therefore, the inverse used for the symmetric network can be reused, with the replacement **w** ↦ (**z, v**).

Using the Sherman–Morrison identity in exactly the same fashion as before, we obtain (see Proposition 3 in Supplementary Methods for the proof):

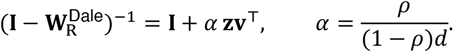

Substituting this expression into Eq. 2 gives (see Proposition 4 in Supplementary Methods for the proof):

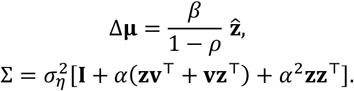

With a decoder 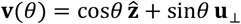, where 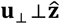 is any unit vector. Algebra identical to the symmetric case yields

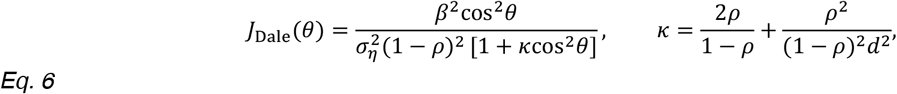

which again peaks at θ = 0^∘^. For the default parameters ρ = 0.4 and *d* ≈ 0.985 we obtain *κ* ≈ 1.79.

Because both rank and leading eigenvalue are preserved, all subsequent analytic results, noise alignment, decoder optimality, and Fisher-ratio shape, carry over unchanged.

### Statistical and Quantification Methods

#### Calculation of stimulus axes and comparison with behavior

To quantify the visual information content that is aligned with the axis of correlated variability, we calculate the correlation of the projection of evoked responses on the axis of correlated variability with the stimulus feature. Where appropriate, we also calculate a feature-specific decoder by training a cross-validated linear regression model and compare the decoder performance with the correlation found above. This common analysis across all our datasets forms the scaffolding of the various tests of our central hypothesis. The details of these analyses are experiment-dependent and are detailed below:

#### Change detection experiments

We first projected the responses for all oriented gratings on the axis of correlated variability. Then, for each starting and change orientation pair, we calculated the performance of a leave-one-out cross-validated linear model for classifying the two orientations. Of all pairs, we selected the most and least aligned pairs and compared the behavioral performance for those pairs (Figure 1C).

#### Curvature estimation experiments

To test whether the information about the visual feature that the monkey uses to guide behavior is the one that has the larger projection on the axis of correlated variability, we first projected all stimulus-evoked responses onto the axis of correlated variability. We then correlated these projections with the curvature feature value or the irrelevant feature value. In other words, we compared the performance of two linear decoders trained on the stimulus-evoked responses projected onto the axis of correlated variability. In each session, we tried various combinations of irrelevant features including orientation, color, surface profile, axial thickness, etc. In supplementary analyses, we also compared the average behavioral performance (one minus the average absolute behavioral error) between the most and least aligned shapes, i.e., the ones that had the highest and lowest decoding performance (Figure S3B-D).

#### Arc manipulation experiments

To test whether behavioral planning-related activity or feature-related activity varies along the axis of correlated variability, we trained two linear decoders on the activity immediately following the onset of the target arc – one to decode the curvature of the visual stimulus (like above) and one to decode the planned saccade. We compared the performance of these decoders with the performance of curvature and saccade decoders trained on the same responses but projected onto the axis of correlated variability (Figure 4D). We did this separately for each shape (although previous results suggest that a shape-general curvature and saccade decoder would also work just as well). We calculated the difference between the decoder prediction accuracies of the curvature decoders and the saccade decoders. We found a larger drop in curvature decoding performance when projected onto the axis of correlated variability (Figure 4E).

#### Curvature-color 2AFC experiments

To test whether learning to use a previously irrelevant feature to guide behavior changes its projection on to the axis of correlated variability, we first calculated the projection of the responses to all stimuli (25 stimuli = 5 color variations and 5 curvature variations) on the axis of correlated variability. In early recording sessions, monkeys 1 and 2 were proficient in using curvature and color only to guide choices, respectively. In later sessions, both monkeys were proficient in using both features. We compared the projection of curvature and color variants of the stimuli on the axis of correlated variability during early and late sessions (Figure 5D-E).

#### Dot direction estimation experiments

To causally test if stimulating neural activity along the axis of correlated variability would have a greater behavioral effect (versus stimulating orthogonal to it), we quantified the neural and behavioral effects of microstimulation. First, we calculated the projection of the effect of microstimulation on the neural responses (measured as the short stimulation-evoked population response vector; details above) on the axis of correlated variability. We performed this analysis separately for both stimulation sites and identified the site with the larger projection. Then, from the long-stim trials, we calculated the size of the behavioral effect of microstimulation (measured as the difference in the slope of the psychometric curve with and without stimulation; details above). We plotted the size of the behavioral effect against the projection on the noise axis calculated above (Figure 6E and Figure S4).

## References

1. M. R. Cohen, A. Kohn, Measuring and interpreting neuronal correlations. Nat. Neurosci. 14, 811–819 (2011).

2. M. R. Cohen, J. H. R. Maunsell, Attention improves performance primarily by reducing interneuronal correlations. Nat. Neurosci. 12, 1594–1600 (2009).

3. G. G. Gregoriou, A. F. Rossi, L. G. Ungerleider, R. Desimone, Lesions of prefrontal cortex reduce attentional modulation of neuronal responses and synchrony in V4. Nat. Neurosci. 17, 1003–1011 (2014).

4. Y. Gu, et al., Perceptual learning reduces interneuronal correlations in macaque visual cortex. Neuron 71, 750–761 (2011).

5. J. L. Herrero, M. A. Gieselmann, M. Sanayei, A. Thiele, Attention-Induced Variance and Noise Correlation Reduction in Macaque V1 Is Mediated by NMDA Receptors. Neuron 78, 729–739 (2013).

6. T. Z. Luo, J. H. R. Maunsell, Neuronal Modulations in Visual Cortex Are Associated with Only One of Multiple Components of Attention. Neuron 86, 1182–1188 (2015).

7. J. P. Mayo, J. H. R. Maunsell, Graded Neuronal Modulations Related to Visual Spatial Attention. J. Neurosci. 36, 5353–5361 (2016).

8. J. F. Mitchell, K. A. Sundberg, J. H. Reynolds, Spatial attention decorrelates intrinsic activity fluctuations in macaque area V4. Neuron 63, 879–888 (2009).

9. A. M. Ni, D. A. Ruff, J. J. Alberts, J. Symmonds, M. R. Cohen, Learning and attention reveal a general relationship between population activity and behavior. Science 359, 463–465 (2018).

10. J. Poort, et al., Learning and attention increase visual response selectivity through distinct mechanisms. Neuron 110, 686-697.e6 (2022).

11. D. A. Ruff, M. R. Cohen, Global cognitive factors modulate correlated response variability between V4 neurons. J. Neurosci. 34, 16408–16416 (2014).

12. D. A. Ruff, M. R. Cohen, Stimulus Dependence of Correlated Variability across Cortical Areas. J. Neurosci. 36, 7546–7556 (2016).

13. D. A. Ruff, J. J. Alberts, M. R. Cohen, Relating normalization to neuronal populations across cortical areas. J. Neurophysiol. 116, 1375–1386 (2016).

14. R. Srinath, D. A. Ruff, M. R. Cohen, Attention improves information flow between neuronal populations without changing the communication subspace. Curr. Biol. 31, 5299-5313.e4 (2021).

15. A. Zénon, R. J. Krauzlis, Attention deficits without cortical neuronal deficits. Nature 489, 434–437 (2012).

16. A. Kohn, R. Coen-Cagli, I. Kanitscheider, A. Pouget, Correlations and neuronal population information. Annu. Rev. Neurosci. 39, 237–256 (2016).

17. H. Nienborg, M. R. Cohen, B. G. Cumming, Decision-related activity in sensory neurons: correlations among neurons and with behavior. Annu. Rev. Neurosci. 35, 463–483 (2012).

18. H. Nienborg, B. Cumming, Correlations between the activity of sensory neurons and behavior: how much do they tell us about a neuron’s causality? Curr. Opin. Neurobiol. 20, 376–381 (2010).

19. Y. Yan, et al., Perceptual training continuously refines neuronal population codes in primary visual cortex. Nat. Neurosci. 17, 1380–1387 (2014).

20. M. R. Cohen, J. H. R. Maunsell, A neuronal population measure of attention predicts behavioral performance on individual trials. J. Neurosci. 30, 15241–15253 (2010).

21. A. M. Ni, B. S. Bowes, D. A. Ruff, M. R. Cohen, Methylphenidate as a causal test of translational and basic neural coding hypotheses. Proc. Natl. Acad. Sci. 119, e2120529119 (2022).

22. M. D. Rosenberg, et al., Methylphenidate Modulates Functional Network Connectivity to Enhance Attention. J. Neurosci. 36, 9547–9557 (2016).

23. R. C. Williamson, et al., Scaling Properties of Dimensionality Reduction for Neural Populations and Network Models. PLOS Comput. Biol. 12, e1005141 (2016).

24. N. C. Rabinowitz, R. L. Goris, M. Cohen, E. P. Simoncelli, Attention stabilizes the shared gain of V4 populations. eLife 4, e08998 (2015).

25. I.-C. Lin, M. Okun, M. Carandini, K. D. Harris, The Nature of Shared Cortical Variability. Neuron 87, 644–656 (2015).

26. R. Moreno-Bote, et al., Information-limiting correlations. Nat. Neurosci. 17, 1410–1417 (2014).

27. M. Kafashan, et al., Scaling of sensory information in large neural populations shows signatures of information-limiting correlations. Nat. Commun. 12, 473 (2021).

28. I. Kanitscheider, R. Coen-Cagli, A. Pouget, Origin of information-limiting noise correlations. Proc. Natl. Acad. Sci. 112, E6973–E6982 (2015).

29. B. B. Averbeck, P. E. Latham, A. Pouget, Neural correlations, population coding and computation. Nat. Rev. Neurosci. 7, 358–366 (2006).

30. D. A. Ruff, A. M. Ni, M. R. Cohen, Cognition as a Window into Neuronal Population Space. Annu. Rev. Neurosci. 41, 77–97 (2018).

31. X. Pitkow, S. Liu, D. E. Angelaki, G. C. DeAngelis, A. Pouget, How Can Single Sensory Neurons Predict Behavior? Neuron 87, 411–423 (2015).

32. O. I. Rumyantsev, et al., Fundamental bounds on the fidelity of sensory cortical coding. Nature 580, 100–105 (2020).

33. D. A. Ruff, M. R. Cohen, Simultaneous multi-area recordings suggest that attention improves performance by reshaping stimulus representations. Nat. Neurosci. 22, 1669–1676 (2019).

34. A. M. Ni, D. A. Ruff, J. J. Alberts, J. Symmonds, M. R. Cohen, Learning and attention reveal a general relationship between population activity and behavior. Science 359, 463–465 (2018).

35. A. M. Ni, C. Huang, B. Doiron, M. R. Cohen, A general decoding strategy explains the relationship between behavior and correlated variability. eLife 11, e67258 (2022).

36. D. A. Ruff, M. R. Cohen, Attention can either increase or decrease spike count correlations in visual cortex. Nat. Neurosci. 17, 1591–1597 (2014).

37. E. Zohary, M. N. Shadlen, W. T. Newsome, Correlated neuronal discharge rate and its implications for psychophysical performance. Nature 370, 140–143 (1994).

38. W. Bair, E. Zohary, W. T. Newsome, Correlated firing in macaque visual area MT: time scales and relationship to behavior. J. Neurosci. Off. J. Soc. Neurosci. 21, 1676–1697 (2001).

39. H. Ko, et al., The emergence of functional microcircuits in visual cortex. Nature 496, 96–100 (2013).

40. H. Ko, T. D. Mrsic-Flogel, S. B. Hofer, Emergence of feature-specific connectivity in cortical microcircuits in the absence of visual experience. J. Neurosci. Off. J. Soc. Neurosci. 34, 9812–9816 (2014).

41. J. C. Eccles, Chemical transmission and Dale’s principle. Prog. Brain Res. 68, 3–13 (1986).

42. A. Chadwick, et al., Learning shapes cortical dynamics to enhance integration of relevant sensory input. Neuron 111, 106-120.e10 (2023).

43. S. B. Hofer, et al., Differential connectivity and response dynamics of excitatory and inhibitory neurons in visual cortex. Nat. Neurosci. 14, 1045–1052 (2011).

44. M. Khosla, A. H. Williams, J. McDermott, N. Kanwisher, Privileged representational axes in biological and artificial neural networks. [Preprint] (2024). Available at: https://www.biorxiv.org/content/10.1101/2024.06.20.599957v1 [Accessed 1 July 2025].

45. M. R. Cohen, W. T. Newsome, Context-Dependent Changes in Functional Circuitry in Visual Area MT. Neuron 60, 162–173 (2008).

46. D. J. Denman, D. Contreras, The structure of pairwise correlation in mouse primary visual cortex reveals functional organization in the absence of an orientation map. Cereb. Cortex N. Y. N 1991 24, 2707–2720 (2014).

47. B. G. Cumming, H. Nienborg, Feedforward and feedback sources of choice probability in neural population responses. Curr. Opin. Neurobiol. 37, 126–132 (2016).

48. X. Huang, S. G. Lisberger, Noise Correlations in Cortical Area MT and Their Potential Impact on Trial-by-Trial Variation in the Direction and Speed of Smooth-Pursuit Eye Movements. J. Neurophysiol. 101, 3012–3030 (2009).

49. M. A. Smith, A. Kohn, Spatial and temporal scales of neuronal correlation in primary visual cortex. J. Neurosci. 28, 12591–12603 (2008).

50. S. G. Solomon, A. Kohn, Moving sensory adaptation beyond suppressive effects in single neurons. Curr. Biol. CB 24, R1012–1022 (2014).

51. A. S. Ecker, et al., Decorrelated neuronal firing in cortical microcircuits. Science 327, 584–587 (2010).

52. B. B. Averbeck, D. Lee, Neural Noise and Movement-Related Codes in the Macaque Supplementary Motor Area. J. Neurosci. 23, 7630–7641 (2003).

53. R. Srinath, M. M. Czarnik, M. R. Cohen, Coordinated Response Modulations Enable Flexible Use of Visual Information. [Preprint] (2024). Available at: https://www.biorxiv.org/content/10.1101/2024.07.10.602774v1 [Accessed 15 July 2024].

54. J. H. R. Maunsell, Neuronal Mechanisms of Visual Attention. Annu. Rev. Vis. Sci. 1, 373–391 (2015).

55. S. Treue, Neural correlates of attention in primate visual cortex. Trends Neurosci. 24, 295–300 (2001).

56. J. H. Reynolds, L. Chelazzi, Attentional Modulation of Visual Processing. Annu. Rev. Neurosci. 27, 611–647 (2004).

57. A. M. Ni, S. Ray, J. H. R. Maunsell, Tuned normalization explains the size of attention modulations. Neuron 73, 803–813 (2012).

58. D. A. Ruff, M. R. Cohen, A normalization model suggests that attention changes the weighting of inputs between visual areas. Proc. Natl. Acad. Sci. 114, E4085–E4094 (2017).

59. K. A. Sundberg, J. F. Mitchell, J. H. Reynolds, Spatial Attention Modulates Center-Surround Interactions in Macaque Visual Area V4. Neuron 61, 952–963 (2009).

60. A. V. Flevaris, S. O. Murray, Feature-based attention modulates surround suppression. J. Vis. 15, 29 (2015).

61. C. Murasugi, C. Salzman, W. Newsome, Microstimulation in visual area MT: effects of varying pulse amplitude and frequency. J. Neurosci. 13, 1719–1729 (1993).

62. C. D. Salzman, K. H. Britten, W. T. Newsome, Cortical microstimulation influences perceptual judgements of motion direction. Nature 346, 174–177 (1990).

63. M. R. Cohen, W. T. Newsome, What electrical microstimulation has revealed about the neural basis of cognition. Curr. Opin. Neurobiol. 14, 169–177 (2004).

64. J. W. Bisley, D. Zaksas, T. Pasternak, Microstimulation of cortical area MT affects performance on a visual working memory task. J. Neurophysiol. 85, 187–196 (2001).

65. M. J. Nichols, W. T. Newsome, Middle Temporal Visual Area Microstimulation Influences Veridical Judgments of Motion Direction. J. Neurosci. 22, 9530–9540 (2002).

66. T. Moore, K. M. Armstrong, Selective gating of visual signals by microstimulation of frontal cortex. Nature 421, 370–373 (2003).

67. D. A. Ruff, S. K. Markman, J. Z. Kim, M. R. Cohen, Linking neural population formatting to function. [Preprint] (2025). Available at: https://www.biorxiv.org/content/10.1101/2025.01.03.631242v1 [Accessed 4 January 2025].

68. C. Xue, L. E. Kramer, M. R. Cohen, Dynamic task-belief is an integral part of decision-making. Neuron 110, 2503-2511.e3 (2022).

69. D. A. Ruff, C. Xue, L. E. Kramer, F. Baqai, M. R. Cohen, Low rank mechanisms underlying flexible visual representations. Proc. Natl. Acad. Sci. 117, 29321–29329 (2020).

70. A. S. Nandy, J. J. Nassi, J. H. Reynolds, Laminar Organization of Attentional Modulation in Macaque Visual Area V4. Neuron 93, 235–246 (2017).

71. C. Stringer, et al., Spontaneous behaviors drive multidimensional, brainwide activity. Science 364, eaav7893 (2019).

72. B. Hayden, J. Gallant, Working Memory and Decision Processes in Visual Area V4. Front. Neurosci. 7 (2013).

73. J. X. Brooks, K. E. Cullen, Predictive Sensing: The Role of Motor Signals in Sensory Processing. Biol. Psychiatry Cogn. Neurosci. Neuroimaging 4, 842–850 (2019).

74. J. C. Martınez-Trujillo, S. Treue, Attentional Modulation Strength in Cortical Area MT Depends on Stimulus Contrast. Neuron 35, 365–370 (2002).

75. R. Srinath, A. M. Ni, C. Marucci, M. R. Cohen, D. H. Brainard, Orthogonal neural representations support perceptual judgements of natural stimuli. [Preprint] (2024). Available at: https://www.biorxiv.org/content/10.1101/2024.02.14.580134v1 [Accessed 19 February 2024].

76. S. Saxena, J. P. Cunningham, Towards the neural population doctrine. Curr. Opin. Neurobiol. 55, 103–111 (2019).

77. R. L. van den Brink, et al., Flexible sensory-motor mapping rules manifest in correlated variability of stimulus and action codes across the brain. Neuron 111, 571-584.e9 (2023).

78. M. T. Kaufman, et al., The Largest Response Component in the Motor Cortex Reflects Movement Timing but Not Movement Type. eNeuro 3 (2016).

79. A. G. Bondy, R. M. Haefner, B. G. Cumming, Feedback determines the structure of correlated variability in primary visual cortex. Nat. Neurosci. 21, 598–606 (2018).

80. T. Kenet, D. Bibitchkov, M. Tsodyks, A. Grinvald, A. Arieli, Spontaneously emerging cortical representations of visual attributes. Nature 425, 954–956 (2003).

81. P. Berkes, G. Orbán, M. Lengyel, J. Fiser, Spontaneous Cortical Activity Reveals Hallmarks of an Optimal Internal Model of the Environment. Science 331, 83–87 (2011).

82. J. Fiser, P. Berkes, G. Orbán, M. Lengyel, Statistically optimal perception and learning: from behavior to neural representations. Trends Cogn. Sci. 14, 119–130 (2010).

83. R. M. Haefner, P. Berkes, J. Fiser, Perceptual Decision-Making as Probabilistic Inference by Neural Sampling. Neuron 90, 649–660 (2016).

84. R. D. Lange, R. M. Haefner, Task-induced neural covariability as a signature of approximate Bayesian learning and inference. PLOS Comput. Biol. 18, e1009557 (2022).

85. G. Orbán, P. Berkes, J. Fiser, M. Lengyel, Neural Variability and Sampling-Based Probabilistic Representations in the Visual Cortex. Neuron 92, 530–543 (2016).

86. R. Echeveste, M. Lengyel, The Redemption of Noise: Inference with Neural Populations. Trends Neurosci. 41, 767–770 (2018).

87. W. J. Ma, J. M. Beck, P. E. Latham, A. Pouget, Bayesian inference with probabilistic population codes. Nat. Neurosci. 9, 1432–1438 (2006).

88. S. A. Moosavi, S. S. R. Hindupur, H. Shimazaki, Population coding under the scale-invariance of high-dimensional noise. [Preprint] (2025). Available at: https://www.biorxiv.org/content/10.1101/2024.08.23.608710v2 [Accessed 22 February 2026].

89. G. Mahuas, T. Buffet, O. Marre, U. Ferrari, T. Mora, Strong, but not weak, noise correlations are beneficial for population coding. [Preprint] (2024). Available at: https://www.biorxiv.org/content/10.1101/2024.06.26.600826v1 [Accessed 22 February 2026].

90. M. H. Histed, A. M. Ni, J. H. R. Maunsell, Insights into cortical mechanisms of behavior from microstimulation experiments. Prog. Neurobiol. 103, 115–130 (2013).

91. A. Szelényi, et al., Intraoperative electrical stimulation in awake craniotomy: methodological aspects of current practice. Neurosurg. Focus 28, E7 (2010).

92. S. N. Flesher, et al., Intracortical microstimulation of human somatosensory cortex. Sci. Transl. Med. 8, 361ra141 (2016).

93. K. Nowik, E. Langwińska-Wośko, P. Skopiński, K. E. Nowik, J. P. Szaflik, Bionic eye review – An update. J. Clin. Neurosci. 78, 8–19 (2020).

94. A. Farnum, G. Pelled, New Vision for Visual Prostheses. Front. Neurosci. 14, 36 (2020).

95. P. R. Roelfsema, Writing to the Mind’s Eye of the Blind. Cell 181, 758–759 (2020).

96. A. L. Anwyl-Irvine, J. Massonnié, A. Flitton, N. Kirkham, J. K. Evershed, Gorilla in our midst: An online behavioral experiment builder. Behav. Res. Methods 52, 388–407 (2020).

97. D. H. Brainard, The Psychophysics Toolbox. Spat. Vis. 10, 433–436 (1997).

