## Supplementary Figures and Appendices for "Guided by Noise: Correlated Variability Channels Task-Relevant Information in Sensory Neurons"

### Supplementary Information

**Appendix A – When do the stimulus response axis and the axis of correlated variability coincide?** **11**

**Appendix B – Decoding optimality along the axis of correlated variability** **14**

**Appendix C – Single-axis Fisher information Changes** **18**

**Appendix D – Optimal encoding for temporal discrimination** **22**

### Supplementary Figures

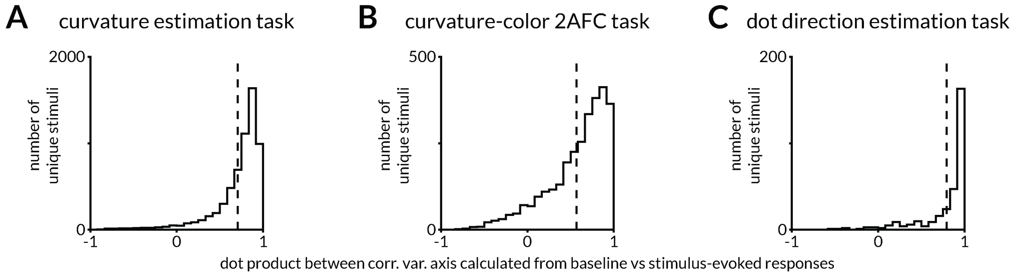

**Figure S1: Axes of correlated variability calculated from baseline period and repeated presentations of the same stimulus are aligned.**

For all three datasets, we calculated the axis of correlated variability in two ways: (1) as the first principal component of the baseline responses (spike rate in a 150-200 ms window after the monkey starts fixating and before the visual stimulus is presented), and (2) as the first principal component of the responses to each unique stimulus. Method 2 would yield as many axes of correlated variability as there were unique visual stimuli. We calculated the dot product of each of these axes with the one calculated with Method 1. The histograms of those dot products are presented in the panels above. Across the three datasets, we found high alignment between the axes measured using the two methods. Even though this analysis shows that the axis of correlated variability is stimulus-agnostic, we measured it using Method 1 across datasets for consistency.

**A.** number of sessions = 124, number of unique stimuli per session = 30-80 (mean 52), number of trials per session = 202-1021 (mean 446), mean dot product = 0.708.

**B.** number of sessions = 118, number of unique stimuli per session = 25, number of trials per session = 121-1466 (mean 593), mean dot product = 0.57.

**C.** number of sessions = 18, number of unique stimuli per session = 10-25 (mean 17.2), number of trials per session = 106-2854 (mean 1199), mean dot product = 0.793.

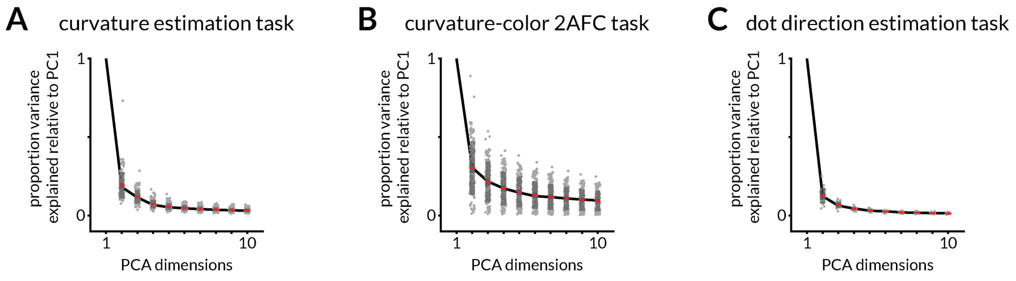

**Figure S2: The first principal component of baseline activity explains a majority of variance across datasets.**

For all three datasets, we show the proportion of variance explained relative to the first principal component. Number of sessions and trials are the same as Figure S1.

**A.** Median percentage of total variance explained by PC1 = 38.6% for monkey 1 and 43.2% for monkey 2. By PC2 = 9.2% for monkey 1 and 5.7% for monkey 2.

**B.** Median percentage of total variance explained by PC1 = 18.8% for monkey 1 and 47.7% for monkey 2. By PC2 = 7.4% for monkey 1 and 7.2% for monkey 2.

**C.** Median percentage of total variance explained by PC1 = 65.7% for monkey 1 and 68.6% for monkey 2. By PC2 = 7.6% for monkey 1 and 10.4% for monkey 2.

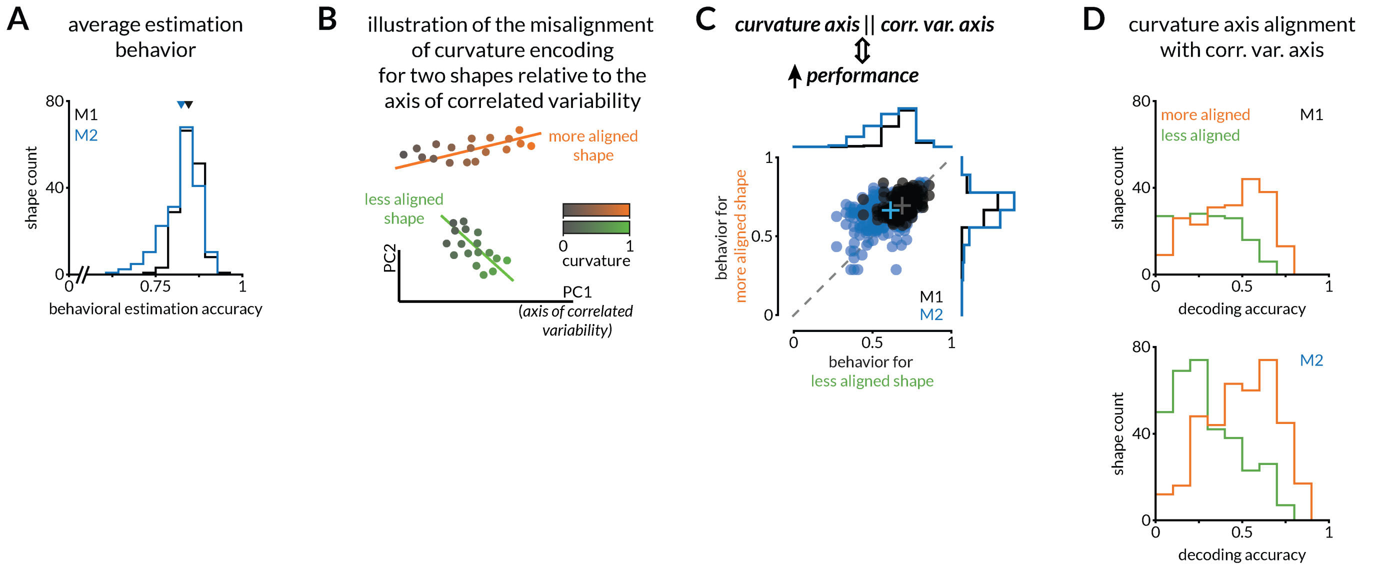

**Figure S3: Behavior summary and additional hypothesis testing related to curvature axis alignment with the axis of correlated variability (Related to Figure 3).**

**A.** Histogram of average estimation behavior (defined as 1-mean absolute difference in stimulus curvature and choice) for each shape tested across 63 sessions for monkey 1 (M1 in black) and 61 sessions for monkey 2 (M2 in blue). Total shapes tested = 199 and 239, and mean $\pm$ SEM = $0.85 \pm 0.03$ and $0.824 \pm 0.054$ for the two monkeys respectively.

**B.** Our hypothesis predicts that performance for a shape with a curvature representation that aligns with the axis of correlated variability is better than one that does not.

**C.** Validation of the prediction in B. Each dot represents behavioral performance for a pair of shapes tested during the same experimental session. Performance was better for the shape that happened to be better aligned to the axis of correlated variability for monkey 2 ($p=1.9 \times 10^{-17}$; n=381 shape pairs; Wilcoxon signed rank test) but not monkey 1 (p=0.45; n=219 shape pairs; Wilcoxon signed rank test). Conventions as in Figure 1C.

**D.** Histograms comparing curvature decoding accuracies along the axis of correlated variability for the more and less aligned shapes in C. For monkey 1 (top), total shape pairs tested = 219, and mean $\pm$ SEM = $0.16 \pm 0.02$ for and $0.42 \pm 0.013$ for the less and more aligned shapes respectively with a KL divergence of 43.83. For monkey 2 (bottom), total shape pairs tested = 381, and mean $\pm$ SEM = $0.245 \pm 0.01$ for and $0.49 \pm 0.01$ for the less and more aligned shapes respectively with a KL divergence of 128.67. The mean difference between the decoding accuracies of the more and less aligned shapes was 0.17 for monkey 1 and 0.223 for monkey 2. One reason for the lack of a significant effect in C for monkey 1 is that the decoding accuracies of pairs of shapes for monkey 1 were more similar to each other (i.e., not considerably better or worse).

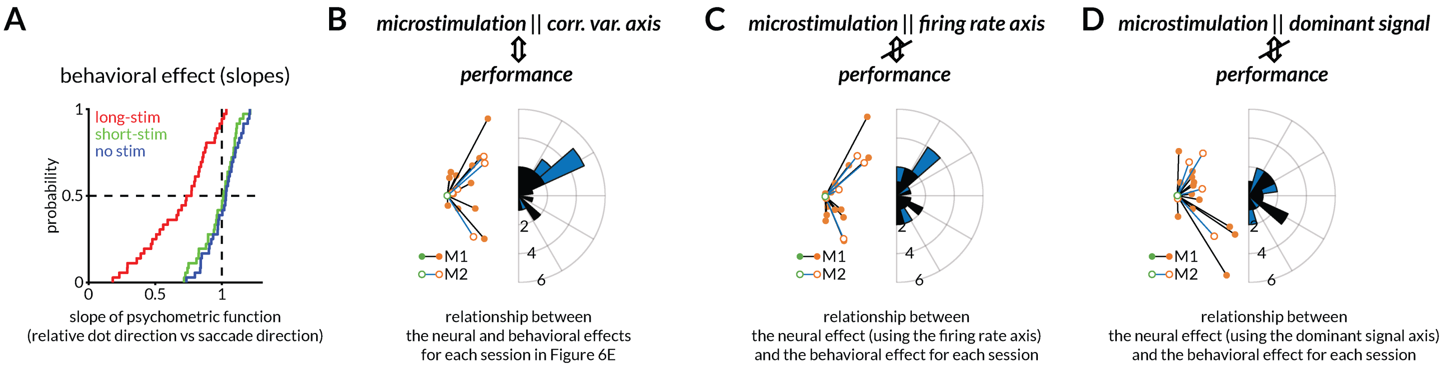

**Figure S4: Behavior summary and quantification and comparison of the effect of electrical microstimulation on behavior (Related to Figure 6)**

**A.** Cumulative distribution of the slope of the psychometric functions for the continuous dot estimation task for the *long-stim* (median 0.753), *short-stim* (median 1.01), and *no stimulation* (median 1.03) conditions. There was no measurable behavioral bias caused by *short-stim*.

**B.** In Figure 6E, each experimental session is depicted by a line connecting the behavioral and neural effect of the two microstimulation sites. To quantify this effect, we centered all the green points (the microstimulation sites with the smaller projection on the axis of correlated variability). By definition, all the orange points (the microstimulation sites with the larger projection on the axis of correlated variability) will be right of the origin. The distribution of the slopes is plotted on the right. The distributions for the two monkeys (M1=black, M2=blue) were significantly skewed (p=0.0037, n=13 and p=0.0307, n=5 for the two monkeys respectively; Rayleigh test for circular non-uniformity).

**C.** We calculated an alternate axis for comparison – the axis that connects the minimum and maximum stimulus-evoked rates for the no electrical stimulation condition. We did the analogous analysis as B with this alternate axis, i.e., we calculated the projection of the effect of microstimulation and compared them with the behavioral effect. The circular distributions of slopes were not significantly non-uniform.

**D.** We calculated another alternate axis – a dominant signal axis defined as the first principal component of the repetition-averaged visually evoked responses. We repeated the same analysis and the circular distributions of slopes were not significantly non-uniform.

### Supplementary Methods (Network Modelling)

#### 1.1 Network dynamics and notation

We modelled a linear rate network of $N = 120$ linear–rate units with activities $\boldsymbol{r}(t) \in \mathbb{R}^N$ that evolve according to

$$\tau\, \dot{\boldsymbol{r}}(t) = -\boldsymbol{r}(t) + \mathbf{G}\mathbf{W}_{\mathrm{R}}\boldsymbol{r}(t) + \mathbf{G}\boldsymbol{w}_{\mathrm{F}}\, s(t) + \boldsymbol{\eta}, \tag{1}$$

where

- $\tau \equiv 1$ is the membrane time-constant;
- $s(t) \in \{0, 1\}$ is a binary stimulus that turns the feed-forward drive on (1) or off (0);
- $\boldsymbol{\eta} \sim \mathcal{N}(\mathbf{0}, \sigma_\eta^2 \mathbf{I})$ is private Gaussian noise with $\sigma_\eta = 1$;
  - Each neuron received an independent and identically distributed Gaussian noise input, ensuring that this component did not introduce shared variability across the population.
- $\mathbf{W}_{\mathrm{R}} \in \mathbb{R}^{N\times N}$ is the recurrent weight matrix;
- $\mathbf{G} = \mathrm{diag}(g_1, \ldots, g_N) \succ 0$ is a static gain matrix;
- $\boldsymbol{w}_{\mathrm{F}} \in \mathbb{R}^N$ is the feed-forward drive vector.

**Steady state.** Whenever the spectral radius of the feedback operator satisfies $\rho(\mathbf{G}\mathbf{W}_{\mathrm{R}}) < 1$, the matrix $\mathbf{I} - \mathbf{G}\mathbf{W}_{\mathrm{R}}$ is invertible, the mean of the stimulus response $\mu_s$ and the noise response $\Sigma$ are

$$\begin{aligned} \boldsymbol{r} \mid s &\sim \mathcal{N}\big(\boldsymbol{\mu}_s,\ \Sigma\big), \\ \boldsymbol{\mu}_s &= (\mathbf{I} - \mathbf{G}\mathbf{W}_{\mathrm{R}})^{-1}\mathbf{G}\boldsymbol{w}_{\mathrm{F}}\, s, &(2)\\ \Sigma &= \sigma_\eta^2\, (\mathbf{I} - \mathbf{G}\mathbf{W}_{\mathrm{R}})^{-1}(\mathbf{I} - \mathbf{G}\mathbf{W}_{\mathrm{R}})^{-\top}. &(3) \end{aligned}$$

We constrained the spectral radius (i.e., the magnitude of the largest eigenvalue) of the recurrent connectivity matrix to be less than one. This condition ensures that recurrent amplification remains bounded and that the network dynamics are stable.

Two expressions will reappear throughout the paper, the difference in stimulus-conditioned means and the Linear Fisher information (unit decoder) are

$$\Delta\boldsymbol{\mu} \coloneqq \boldsymbol{\mu}_1 - \boldsymbol{\mu}_0 = (\mathbf{I} - \mathbf{G}\mathbf{W}_{\mathrm{R}})^{-1}\mathbf{G}\boldsymbol{w}_{\mathrm{F}}, \tag{4a}$$

$$J(\boldsymbol{v}) \coloneqq \frac{\big(\boldsymbol{v}^\top \Delta\boldsymbol{\mu}\big)^2}{\boldsymbol{v}^\top \Sigma\, \boldsymbol{v}}, \qquad \|\boldsymbol{v}\| = 1, \tag{4b}$$

where $J(\boldsymbol{v})$ is the linear Fisher ratio for a one-dimensional decoder $\boldsymbol{v}$.

Unless noted otherwise, we use $N = 120$, $\sigma_\eta = 1$, $\tau = 1$ and $\mathbf{G} = \mathbf{I}$ for both (Sect. 1.2) and (Sect. 1.3).

#### 1.2 Rank-one symmetric network

We are first going to start with a very simple network, where

**Definition**

$$\boldsymbol{w}_{\mathrm{F}} = \beta\, \boldsymbol{w}, \qquad \|\boldsymbol{w}\| = 1, \qquad \beta > 0, \tag{5}$$

$$\mathbf{W}_{\mathrm{R}} = \rho\, \boldsymbol{w}\boldsymbol{w}^{\top}, \qquad 0 < \rho < 1. \tag{6}$$

Only the first unit is stimulated, hence $\boldsymbol{w} = (1, 0, \ldots, 0)^{\top}$ there; the algebra below holds for an arbitrary unit vector $\boldsymbol{w}$. The reason we choose $\boldsymbol{w} = (1, 0, \ldots, 0)^{\top}$ here is to make the feedforward and recurrent the simplest rank-one structure. In the Dale's law compliant network section we will show the more realistic version that we implemented to produce Figure 2.

**Steady-state statistics**

**Proposition 1** (Rank-one inverse)**.** *Let* $\boldsymbol{w} \in \mathbb{R}^N$ *be a unit vector* ($\|\boldsymbol{w}\| = 1$) *and define the rank-one matrix* $\mathbf{W}_{\mathrm{R}} = \rho\, \boldsymbol{w}\boldsymbol{w}^{\top}$ *with* $0 < \rho < 1$*. Then*

$$(\mathbf{I} - \mathbf{W}_{\mathrm{R}})^{-1} \;=\; \mathbf{I} + \frac{\rho}{1-\rho}\, \boldsymbol{w}\boldsymbol{w}^{\top}.$$

*Proof.* We use the classical *Sherman–Morrison identity*, which gives the inverse of a rank-one perturbation of an invertible matrix:

$$(\mathbf{A} - \boldsymbol{u}\boldsymbol{v}^{\top})^{-1} \;=\; \mathbf{A}^{-1} \;+\; \frac{\mathbf{A}^{-1}\boldsymbol{u}\boldsymbol{v}^{\top}\mathbf{A}^{-1}}{1 - \boldsymbol{v}^{\top}\mathbf{A}^{-1}\boldsymbol{u}}, \qquad \text{provided } 1 - \boldsymbol{v}^{\top}\mathbf{A}^{-1}\boldsymbol{u} \neq 0.$$

Take $\mathbf{A} = \mathbf{I}$, $\boldsymbol{u} = \boldsymbol{v} = \sqrt{\rho}\, \boldsymbol{w}$; then $\mathbf{A}^{-1} = \mathbf{I}$ and $\boldsymbol{v}^{\top}\mathbf{A}^{-1}\boldsymbol{u} = (\sqrt{\rho}\, \boldsymbol{w})^{\top}(\sqrt{\rho}\, \boldsymbol{w}) = \rho$. Hence

$$(\mathbf{I} - \rho\, \boldsymbol{w}\boldsymbol{w}^{\top})^{-1} = \mathbf{I} + \frac{\rho\, \boldsymbol{w}\boldsymbol{w}^{\top}}{1-\rho},$$

which is exactly the claimed formula.

For completeness, we can verify it directly by multiplication:

$$\begin{aligned}
&\Big(\mathbf{I} - \rho\, \boldsymbol{w}\boldsymbol{w}^{\top}\Big)\Big(\mathbf{I} + \tfrac{\rho}{1-\rho}\, \boldsymbol{w}\boldsymbol{w}^{\top}\Big) \\
&= \mathbf{I} + \frac{\rho}{1-\rho}\, \boldsymbol{w}\boldsymbol{w}^{\top} - \rho\, \boldsymbol{w}\boldsymbol{w}^{\top} - \frac{\rho^2}{1-\rho}\, \boldsymbol{w}\, \underbrace{(\boldsymbol{w}^{\top}\boldsymbol{w})}_{=1}\, \boldsymbol{w}^{\top} \\
&= \mathbf{I} + \big[\tfrac{\rho}{1-\rho} - \rho - \tfrac{\rho^2}{1-\rho}\big]\boldsymbol{w}\boldsymbol{w}^{\top} = \mathbf{I}.
\end{aligned}$$

The two square brackets cancel because $\frac{\rho}{1-\rho} - \rho - \frac{\rho^2}{1-\rho} = 0$. Therefore, the proposed matrix is indeed the inverse of $\mathbf{I} - \mathbf{W}_{\mathrm{R}}$. □

**Proposition 2** (Symmetric model statistics)**.** *For the rank-one recurrent matrix* $\mathbf{W}_{\mathrm{R}} = \rho\, \boldsymbol{w}\boldsymbol{w}^{\top}$ *with* $0 < \rho < 1$ *and* $\|\boldsymbol{w}\| = 1$ *(see* (6)*), and for* $\boldsymbol{w}_{\mathrm{F}} = \beta\, \boldsymbol{w}$ *used, the quantities in* (4) *are*

$$\Delta\boldsymbol{\mu} = \frac{\beta}{1-\rho}\, \boldsymbol{w}, \tag{7}$$

$$\Sigma = \sigma_\eta^2 \Big[\mathbf{I} + \frac{2\rho - \rho^2}{(1-\rho)^2}\, \boldsymbol{w}\boldsymbol{w}^{\top}\Big]. \tag{8}$$

*Proof.* Lemma 1 gives

$$(\mathbf{I} - \mathbf{W}_{\mathrm{R}})^{-1} = \mathbf{I} + \alpha\, \boldsymbol{w}\boldsymbol{w}^{\top}, \qquad \alpha := \frac{\rho}{1-\rho}. \tag{A}$$

Using definition (4a) with $\boldsymbol{w}_{\mathrm{F}} = \beta\,\boldsymbol{w}$ and (A),

$$\Delta\boldsymbol{\mu} = (\mathbf{I} - \mathbf{W}_{\mathrm{R}})^{-1}\boldsymbol{w}_{\mathrm{F}} = \big(\mathbf{I} + \alpha\,\boldsymbol{w}\boldsymbol{w}^\top\big)\,\beta\boldsymbol{w}.$$

Because $(\boldsymbol{w}\boldsymbol{w}^\top)\boldsymbol{w} = (\boldsymbol{w}^\top\boldsymbol{w})\boldsymbol{w} = \boldsymbol{w}$,

$$\Delta\boldsymbol{\mu} = \beta\big(1+\alpha\big)\boldsymbol{w} = \beta\Big(1 + \frac{\rho}{1-\rho}\Big)\boldsymbol{w} = \frac{\beta}{1-\rho}\,\boldsymbol{w},$$

which is Eq. (7).

Definition (3) (with $\mathbf{W}_{\mathrm{R}}^\top = \mathbf{W}_{\mathrm{R}}$ because the matrix is symmetric) gives

$$\Sigma = \sigma_\eta^2\,\big(\mathbf{I} - \mathbf{W}_{\mathrm{R}}\big)^{-2} = \sigma_\eta^2\,\big[\mathbf{I} + \alpha\,\boldsymbol{w}\boldsymbol{w}^\top\big]^2.$$

Expand the square explicitly:

$$(\mathbf{I} + \alpha\,\boldsymbol{w}\boldsymbol{w}^\top)^2 = \mathbf{I} + 2\alpha\,\boldsymbol{w}\boldsymbol{w}^\top + \alpha^2\big(\boldsymbol{w}\boldsymbol{w}^\top\big)^2.$$

Since $(\boldsymbol{w}\boldsymbol{w}^\top)^2 = \boldsymbol{w}(\boldsymbol{w}^\top\boldsymbol{w})\boldsymbol{w}^\top = \boldsymbol{w}\boldsymbol{w}^\top$ (because $\|\boldsymbol{w}\| = 1$),

$$(\mathbf{I} + \alpha\,\boldsymbol{w}\boldsymbol{w}^\top)^2 = \mathbf{I} + (2\alpha + \alpha^2)\,\boldsymbol{w}\boldsymbol{w}^\top.$$

Insert $\alpha = \rho/(1-\rho)$:

$$2\alpha + \alpha^2 = \frac{2\rho}{1-\rho} + \frac{\rho^2}{(1-\rho)^2} = \frac{2\rho - \rho^2}{(1-\rho)^2}.$$

Hence

$$\Sigma = \sigma_\eta^2\Big[\mathbf{I} + \frac{2\rho-\rho^2}{(1-\rho)^2}\,\boldsymbol{w}\boldsymbol{w}^\top\Big],$$

which is Eq. (8). □

##### Fisher ratio along a 1-d decoder

For a decoder $\boldsymbol{v}(\theta) = \cos\theta\,\boldsymbol{w} + \sin\theta\,\boldsymbol{u}_\perp$ with $\boldsymbol{u}_\perp \perp \boldsymbol{w}$. Using Prop. 2 in (4b) gives for our symmetric matrix $W_{\mathrm{R}}$

$$J_{\mathrm{sym}}(\theta) = \frac{\beta^2\cos^2\theta}{\sigma_\eta^2\big[\cos^2\theta + (1-\rho)^2\sin^2\theta\big]}. \tag{9}$$

$J_{\mathrm{sym}}(\theta)$ is maximized at $\theta = 0$, in agreement with the simulations. The general case of decoding optimality along the axis of correlated variability is shown in Appendix B.

#### 1.3 Rank-one Dale-compliant network

This network, used to generate Figure 2, obeys Dale's law by assigning exclusive excitatory (+) or inhibitory (−) signs to each *row* of the recurrent weight matrix. All other hyper-parameters ($N = 120$, $\sigma_\eta = 1$, $\mathbf{G} = \mathbf{I}$) match the symmetric network of Sect. 1.2.

##### Magnitude profile and sign vector

1. **Gaussian magnitude.** On the grid $x_i = -1 + 2(i-1)/(N-1)$ $(i = 1,\ldots,N)$ set $m_i = \exp(-\kappa x_i^2)$ with $\kappa = 4$. Normalize: $\boldsymbol{m}_{\mathrm{mag}} := (m_1,\ldots,m_N)^\top$, $\boldsymbol{m} := \boldsymbol{m}_{\mathrm{mag}}/\|\boldsymbol{m}_{\mathrm{mag}}\|$, $\|\boldsymbol{m}\| = 1$.
2. **Dale signs.** Rows 1 : 96 (E) are +; rows 97 : 120 (I) are −. Define $\boldsymbol{z} = \mathrm{sgn} \odot \boldsymbol{m}$ and the alignment coefficient $d := \boldsymbol{m}^\top\boldsymbol{z} \in (0,1)$, $d \simeq 0.985$.
3. **Feedforward drive.** Stimulate along the Perron vector (to be shown below): $\boldsymbol{w}_{\mathrm{F}} = \beta\,\hat{\boldsymbol{z}}$, $\hat{\boldsymbol{z}} := \boldsymbol{z}/\|\boldsymbol{z}\| = \boldsymbol{z}$.

**Recurrent matrix and its spectrum**

$$\boxed{\mathbf{W}_{\mathrm{R}}^{\mathrm{Dale}} := \frac{\rho}{d}\,\boldsymbol{z}\boldsymbol{m}^{\top}}, \qquad 0 < \rho < 1. \tag{10}$$

**Why this construction?**

- Right-multiplying by $\boldsymbol{z}$ gives $\mathbf{W}_{\mathrm{R}}^{\mathrm{Dale}}\boldsymbol{z} = \rho\,\boldsymbol{z}$, so $\boldsymbol{z}$ is the Perron (slow/noise) eigenvector with eigenvalue $\rho$, matching the spectral radius of Sect. 1.2.
- Being rank-one keeps the algebra tractable while rotating the Perron vector away from the pure-E direction $\boldsymbol{m}$.

**Eigen-spectrum – detailed derivation.** We prove that the Dale matrix $\mathbf{W}_{\mathrm{R}}^{\mathrm{Dale}} := \frac{\rho}{d}\,\boldsymbol{z}\boldsymbol{m}^{\top}$ has a single non-zero eigenvalue $\rho$ and is rank 1.

**1. $\boldsymbol{z}$ is an eigenvector.**

$$\mathbf{W}_{\mathrm{R}}^{\mathrm{Dale}}\boldsymbol{z} = \frac{\rho}{d}\,\boldsymbol{z}\big(\boldsymbol{m}^{\top}\boldsymbol{z}\big) = \frac{\rho}{d}\,d\,\boldsymbol{z} = \rho\,\boldsymbol{z}.$$

Hence $\lambda_1 = \rho$ with (right) eigenvector $\boldsymbol{z}$.

**2. The image of $\mathbf{W}_{\mathrm{R}}^{\mathrm{Dale}}$ is one-dimensional.** Take an arbitrary vector $\boldsymbol{x} \in \mathbb{R}^N$ and write the scalar $\alpha := \boldsymbol{m}^{\top}\boldsymbol{x} \in \mathbb{R}$. Then

$$\mathbf{W}_{\mathrm{R}}^{\mathrm{Dale}}\boldsymbol{x} = \frac{\rho}{d}\,\boldsymbol{z}\alpha = \big(\tfrac{\rho\alpha}{d}\big)\boldsymbol{z} \;\in\; \operatorname{span}\{\boldsymbol{z}\}.$$

Thus, every image vector is a multiple of $\boldsymbol{z}$; conversely $\boldsymbol{z}$ itself lies in the image because $\mathbf{W}_{\mathrm{R}}^{\mathrm{Dale}}\boldsymbol{z} = \rho\boldsymbol{z} \neq \mathbf{0}$. Therefore,

$$\operatorname{Im}\big(\mathbf{W}_{\mathrm{R}}^{\mathrm{Dale}}\big) = \operatorname{span}\{\boldsymbol{z}\},$$

so the image has dimension 1.

**3. Rank and the zero eigenvalue multiplicity.** Since the rank of a matrix equals the dimension of its image, $\operatorname{rank}\big(\mathbf{W}_{\mathrm{R}}^{\mathrm{Dale}}\big) = 1$. For an $N \times N$ matrix of rank 1 there are $N - 1$ zero eigenvalues (counted with algebraic multiplicity). Together with $\lambda_1 = \rho$ from 1 we have the full spectrum

$$\operatorname{spec}\big(\mathbf{W}_{\mathrm{R}}^{\mathrm{Dale}}\big) = \{\rho,\, 0, \ldots, 0\}, \qquad \rho \text{ (once)},\ 0\ (N-1 \text{ times}).$$

**Interpretation.** Relative to the symmetric model (Sect. 1.2) the non-zero eigenvalue is unchanged ($\rho$), but its eigenvector rotates from $\boldsymbol{m}$ to $\boldsymbol{z}$, reflecting the E–I sign structure imposed by Dale's law while preserving overall stability ($\rho < 1$).

**Steady-state statistics**

**Proposition 3** (Inverse of $\mathbf{I} - \mathbf{W}_{\mathrm{R}}^{\mathrm{Dale}}$)**.**

$$(\mathbf{I} - \mathbf{W}_{\mathrm{R}}^{\mathrm{Dale}})^{-1} = \mathbf{I} + \alpha\,\boldsymbol{z}\boldsymbol{m}^{\top}, \qquad \alpha := \frac{\rho}{(1-\rho)d}.$$

*Proof.* Set $u = \sqrt{\rho/d}\,\boldsymbol{z},\ v = \sqrt{\rho/d}\,\boldsymbol{m}$. The Sherman–Morrison identity for rank-one updates, $(\mathbf{I} - uv^{\top})^{-1} = \mathbf{I} + \frac{uv^{\top}}{1 - v^{\top}u}$, gives

$$(\mathbf{I} - \mathbf{W}_{\mathrm{R}}^{\mathrm{Dale}})^{-1} = \mathbf{I} + \frac{(\rho/d)\,\boldsymbol{z}\boldsymbol{m}^{\top}}{1 - (\rho/d)(\boldsymbol{m}^{\top}\boldsymbol{z})} = \mathbf{I} + \alpha\,\boldsymbol{z}\boldsymbol{m}^{\top}$$

because $\boldsymbol{m}^{\top}\boldsymbol{z} = d$. A direct multiplication check confirms the inverse. □

**Proposition 4** (Dale model statistics)**.** *Using Lemma 3 together with the definitions in (2)–(4),*

$$\Delta\boldsymbol{\mu} = \frac{\beta}{1-\rho}\,\hat{\boldsymbol{z}}, \tag{11}$$

$$\Sigma = \sigma_\eta^2\Big[\mathbf{I} + \alpha\big(\boldsymbol{z}\boldsymbol{m}^\top + \boldsymbol{m}\boldsymbol{z}^\top\big) + \alpha^2\boldsymbol{z}\boldsymbol{z}^\top\Big]. \tag{12}$$

*Proof. Mean shift.* With $\boldsymbol{w}_{\mathrm{F}} = \beta\,\hat{\boldsymbol{z}}$,

$$\Delta\boldsymbol{\mu} = (\mathbf{I} - \mathbf{W}_{\mathrm{R}}^{\mathrm{Dale}})^{-1}\boldsymbol{w}_{\mathrm{F}} = (\mathbf{I} + \alpha\,\boldsymbol{z}\boldsymbol{m}^\top)\beta\,\boldsymbol{z} = \beta\big(1 + \alpha\,(\boldsymbol{m}^\top\boldsymbol{z})\big)\boldsymbol{z} = \beta(1+\alpha d)\boldsymbol{z} = \frac{\beta}{1-\rho}\,\hat{\boldsymbol{z}},$$

since $(\boldsymbol{z}\boldsymbol{m}^\top)\boldsymbol{z} = (\boldsymbol{m}^\top\boldsymbol{z})\boldsymbol{z} = d\,\boldsymbol{z}$ and

$$1 + \alpha d = 1 + \frac{\rho}{(1-\rho)d}\,d = 1 + \frac{\rho}{1-\rho} = \frac{1}{1-\rho}.$$

*Noise covariance.* Eq. (3) yields

$$\Sigma = \sigma_\eta^2\,(\mathbf{I} + \alpha\,\boldsymbol{z}\boldsymbol{m}^\top)(\mathbf{I} + \alpha\,\boldsymbol{m}\boldsymbol{z}^\top).$$

Expanding the product yields:

$$\Sigma = \sigma_\eta^2\Big[\mathbf{I} + \alpha(\boldsymbol{z}\boldsymbol{m}^\top + \boldsymbol{m}\boldsymbol{z}^\top) + \alpha^2(\boldsymbol{z}\boldsymbol{m}^\top)(\boldsymbol{m}\boldsymbol{z}^\top)\Big].$$

Since $(\boldsymbol{z}\boldsymbol{m}^\top)(\boldsymbol{m}\boldsymbol{z}^\top) = (\boldsymbol{m}^\top\boldsymbol{m})\,\boldsymbol{z}\boldsymbol{z}^\top = \boldsymbol{z}\boldsymbol{z}^\top$, we obtain Eq. (12). □

##### Fisher ratio

For a decoder $\boldsymbol{v}(\theta) = \cos\theta\,\hat{\boldsymbol{z}} + \sin\theta\,\boldsymbol{u}_\perp$ choose $\boldsymbol{u}_\perp$ to be a unit vector satisfying

$$\boldsymbol{u}_\perp \perp \hat{\boldsymbol{z}} \quad \text{and} \quad \boldsymbol{u}_\perp \perp \boldsymbol{m}.$$

(This is always possible when $N \geq 3$.) Using Prop. 4 we obtain

$$S_{\mathrm{Dale}}(\theta) = \frac{\beta^2}{(1-\rho)^2}\cos^2\theta,$$

$$N_{\mathrm{Dale}}(\theta) = \sigma_\eta^2\big[1 + \kappa\cos^2\theta\big], \qquad \kappa := \frac{2\rho}{1-\rho} + \frac{\rho^2}{(1-\rho)^2 d^2}, \tag{13}$$

$$J_{\mathrm{Dale}}(\theta) = \frac{\beta^2\cos^2\theta}{\sigma_\eta^2(1-\rho)^2[1+\kappa\cos^2\theta]}. \tag{14}$$

With the default parameters $\rho = 0.4$, $d \simeq 0.98456$ one obtains $\kappa \simeq 1.792$.

### Appendix A – When do the stimulus response axis and the axis of correlated variability coincide?

In this section, we give the conditions under which the stimulus response axis aligns with the dominant noise (slow-mode) axis.

#### A.1 Network and notation

Consider a population of $N$ linear rate units obeying

$$\tau \, \dot{\boldsymbol{r}}(t) = -\boldsymbol{r}(t) + \mathbf{G}\mathbf{W}_{\mathrm{R}}\boldsymbol{r}(t) + \mathbf{G}\boldsymbol{w}_F \, s(t) + \boldsymbol{\eta}, \tag{15}$$

where the symbols in Table S1 retain the meanings used in the main text.

Table S1: Symbols used in Appendix A.

| Symbol | Description |
|---|---|
| $\mathbf{W}_{\mathrm{R}} \in \mathbb{R}^{N\times N}$ | recurrent weight matrix, $\rho(\mathbf{G}\mathbf{W}_{\mathrm{R}}) < 1$ |
| $\mathbf{G} = \mathrm{diag}(g_1, \ldots, g_N) \succ 0$ | static neuronal gains |
| $\boldsymbol{w}_F \in \mathbb{R}^N$ | feed-forward drive vector |
| $s(t) \in \{0, 1\}$ | binary stimulus |
| $\boldsymbol{\eta} \sim \mathcal{N}(\mathbf{0}, \sigma_\eta^2 \mathbf{I})$ | private Gaussian noise |
| $\tau$ | neuronal time-constant (set to 1) |

Equation (15) can be rewritten as

$$\tau \, \dot{\boldsymbol{r}}(t) = -\boldsymbol{r}(t) + \mathbf{G}\mathbf{W}_{\mathrm{R}}\boldsymbol{r}(t) + \mathbf{G}\mathbf{W}_{\mathrm{F}} s(t) + \boldsymbol{\eta}, \qquad \mathbf{W}_{\mathrm{F}} \coloneqq \boldsymbol{w}_F \, \boldsymbol{e}^\top. \tag{16}$$

**Feedback operator.** We assume $\mathbf{A} \coloneqq \mathbf{G}\mathbf{W}_{\mathrm{R}}$ is a real-symmetric matrix, so it is diagonalized by an orthonormal set $\{\boldsymbol{u}_i\}_{i=1}^N$:

$$\mathbf{A}\,\boldsymbol{u}_i = \lambda_i \boldsymbol{u}_i, \qquad 1 > \lambda_1 > \lambda_2 \geq \cdots \geq \lambda_N > -1.$$

The dominant eigen-vector $\boldsymbol{u}_1$ defines the principal direction of internally generated correlated variability.

##### Stimulus Response Axis

Setting $\dot{\boldsymbol{r}} = \mathbf{0}$ in Eq. (15) and conditioning on $s \in \{0, 1\}$ gives (after averaging over noise realizations):

$$\boldsymbol{\mu}_s = (\mathbf{I} - \mathbf{A})^{-1}\mathbf{G}\boldsymbol{w}_F \, s. \tag{17}$$

We denote the difference in stimulus-conditioned means as

$$\Delta\boldsymbol{\mu} \coloneqq \boldsymbol{\mu}_1 - \boldsymbol{\mu}_0 = (\mathbf{I} - \mathbf{A})^{-1}\mathbf{G}\boldsymbol{w}_F. \tag{18}$$

The stimulus response axis is the unit vector

$$\boldsymbol{u}_{\mathrm{stim}} \coloneqq \frac{\Delta\boldsymbol{\mu}}{\|\Delta\boldsymbol{\mu}\|}. \tag{19}$$

#### A.2 Spectral expansion of the stimulus response axis

We first expand $\mathbf{G}\boldsymbol{w}_F$ in the eigenbasis,

$$\mathbf{G}\boldsymbol{w}_F = \sum_{i=1}^{N} c_i\, \boldsymbol{u}_i, \qquad c_i := \boldsymbol{u}_i^\top \mathbf{G}\boldsymbol{w}_F. \tag{20}$$

We next use the Neumann series to rewrite $(\mathbf{I}-\mathbf{A})^{-1}$ as:

$$(\mathbf{I}-\mathbf{A})^{-1} = \sum_{k=0}^{\infty} \mathbf{A}^k, \tag{21}$$

where we assume that $\rho(\mathbf{A}) < 1$ to ensure convergence. We then substitute Eqs. (20) and (21) into Eq. (18) to give:

$$\begin{aligned}
\Delta\boldsymbol{\mu} &= (\mathbf{I}-\mathbf{A})^{-1}\mathbf{G}\boldsymbol{w}_F \\
&= \sum_{k=0}^{\infty} \mathbf{A}^k \Big(\sum_{i=1}^{N} c_i\, \boldsymbol{u}_i\Big) \\
&= \sum_{i=1}^{N} c_i \Big(\sum_{k=0}^{\infty} \lambda_i^k\Big)\boldsymbol{u}_i \qquad (\mathbf{A}^k\boldsymbol{u}_i = \lambda_i^k \boldsymbol{u}_i) \\
&= \sum_{i=1}^{N} \frac{c_i}{1-\lambda_i}\, \boldsymbol{u}_i
\end{aligned}$$

where the geometric series converges because $|\lambda_i| < 1$ for all $i$.

Therefore, we obtain

$$\Delta\boldsymbol{\mu} = \sum_{i=1}^{N} \frac{c_i}{1-\lambda_i}\, \boldsymbol{u}_i. \tag{22}$$

The factor $(1-\lambda_i)^{-1}$ preferentially amplifies slower modes ($\lambda_i \approx 1$).

#### A.3 Necessary and sufficient condition for alignment

**Proposition 5** (Alignment criterion)**.** *With $c_i := \boldsymbol{u}_i^\top \mathbf{G}\boldsymbol{w}_F$ from (20), the following two statements are equivalent:*

$$\underbrace{\big(\boldsymbol{u}_{stim} \parallel \boldsymbol{u}_1\big)}_{\textit{axes coincide}} \iff \underbrace{\big(\mathbf{G}\boldsymbol{w}_F = c_1\boldsymbol{u}_1,\ c_1 \neq 0\big)}_{\textit{drive on noise mode}}.$$

*Equivalently, the axes coincide* iff *$c_i = 0$ for every $i > 1$ while $c_1 \neq 0$.*

*Proof.* Write the stimulus-conditioned mean difference in the eigenbasis (22):

$$\Delta\boldsymbol{\mu} = \sum_{i=1}^{N} \frac{c_i}{1-\lambda_i}\, \boldsymbol{u}_i, \qquad |\lambda_i| < 1.$$

($\Longrightarrow$) **Necessity.** Assume the axes are collinear: $\boldsymbol{u}_{\text{stim}} = \pm\boldsymbol{u}_1 \implies \Delta\boldsymbol{\mu} = \alpha\boldsymbol{u}_1$ for some $\alpha \neq 0$ (the case $\alpha = 0$ would leave $\boldsymbol{u}_{\text{stim}}$ undefined). Project both sides onto each $\boldsymbol{u}_i$:

$$\boldsymbol{u}_i^\top \Delta\boldsymbol{\mu} = \frac{c_i}{1-\lambda_i} = \begin{cases} \alpha & i = 1, \\ 0 & i > 1. \end{cases}$$

Because $1-\lambda_i \neq 0$ for every $i$, we obtain $c_i = 0$ for all $i > 1$ and, in particular, $c_1 = \alpha(1-\lambda_1) \neq 0$. Hence $\mathbf{G}\boldsymbol{w}_F = c_1\boldsymbol{u}_1$.

($\Longleftarrow$) **Sufficiency.** Conversely, assume $\mathbf{G}\boldsymbol{w}_F = c_1\boldsymbol{u}_1$ with $c_1 \neq 0$, so $c_i = 0$ for $i > 1$. Substituting into (22) collapses the sum to

$$\Delta\boldsymbol{\mu} = \frac{c_1}{1-\lambda_1}\,\boldsymbol{u}_1 = \beta\boldsymbol{u}_1, \qquad \beta \neq 0,$$

whence $\boldsymbol{u}_{\text{stim}} = \pm\boldsymbol{u}_1$. Thus, the axes coincide. □

In conclusion, we showed that stimulus–recurrent alignment is not guaranteed; it requires a specific relation between the feedforward drive and the recurrent connectivity.

### Appendix B – Decoding optimality along the axis of correlated variability

Here, we prove that when the stimulus response axis and the axis of correlated variability align (see Appendix A), then the Fisher Ratio for any one-dimensional decoder is maximized. We assume the alignment condition:

$$\Delta\boldsymbol{\mu} \;=\; a\,\boldsymbol{u}_1, \qquad a \;=\; \|\Delta\boldsymbol{\mu}\| > 0.$$

In effect, we assume that the stimulus-conditioned mean shift is *already* aligned with the dominant noise eigenvector $\boldsymbol{u}_1$ of $\mathbf{A} = \mathbf{G}\mathbf{W}_{\mathrm{R}}$. Under this condition, we can prove that the Fisher ratio for any one-dimensional decoder is maximized when the read-out aligns with $\boldsymbol{u}_1$.

#### B.1 Fisher ratio and eigenbasis

For a unit-length 1-d decoder $\boldsymbol{v} \in \mathbb{R}^N$ we define the linear Fisher ratio as:

$$J(\boldsymbol{v}) \;=\; \frac{\left(\boldsymbol{v}^\top \Delta\boldsymbol{\mu}\right)^2}{\boldsymbol{v}^\top \Sigma \boldsymbol{v}}, \qquad \Sigma \;=\; \sigma_\eta^2\,(\mathbf{I}-\mathbf{A})^{-1}(\mathbf{I}-\mathbf{A})^{-\top}. \tag{23}$$

Because the response variance $\Sigma$ shares the eigenbasis of $\mathbf{A}$ we have:

$$\Sigma = \sigma_\eta^2\ \mathrm{diag}\big(d_1, \ldots, d_N\big), \quad d_i \;=\; \frac{1}{(1-\lambda_i)^2}, \qquad d_1 > d_2 \geq \cdots \geq d_N > 0,$$

where the ordering $d_1 > d_2$ follows from $\lambda_1 > \lambda_2$. We introduce an orthonormal vector $\boldsymbol{u}_\perp$ that spans the subspace orthogonal to $\boldsymbol{u}_1$. Any unit decoder can be written as:

$$\boldsymbol{v}(\theta) \;=\; \cos\theta\,\boldsymbol{u}_1 \;+\; \sin\theta\,\boldsymbol{u}_\perp, \qquad \theta \in \left[-\tfrac{\pi}{2}, \tfrac{\pi}{2}\right]. \tag{24}$$

Here, $\theta$ is the angle between the decoder axis and $\boldsymbol{u}_1$ axis.

#### B.2 Closed-form Fisher ratio

Combining Eq. (24) and $\Delta\boldsymbol{\mu} = a\boldsymbol{u}_1$ yields:

$$\left(\boldsymbol{v}^\top \Delta\boldsymbol{\mu}\right)^2 = a^2\cos^2\theta \equiv \mathcal{N}(\theta), \tag{25}$$

$$\boldsymbol{v}^\top \Sigma \boldsymbol{v} = \sigma_\eta^2\big(d_1\cos^2\theta + d_2\sin^2\theta\big) \equiv \mathcal{D}(\theta), \tag{26}$$

so that the Fisher ratio is:

$$J(\theta) = \frac{a^2\cos^2\theta}{\sigma_\eta^2\big(d_1\cos^2\theta + d_2\sin^2\theta\big)} = \frac{\mathcal{N}(\theta)}{\mathcal{D}(\theta)}. \tag{27}$$

#### B.3 First derivative

**Step 1: Derivatives Basics.**

$$\mathcal{N}'(\theta) = a^2 \frac{\mathrm{d}}{\mathrm{d}\theta}\big(\cos^2\theta\big) = a^2 \cdot 2\cos\theta(-\sin\theta) = -2a^2\cos\theta\sin\theta,$$

$$\begin{aligned}\mathcal{D}'(\theta) &= \sigma_\eta^2 \frac{\mathrm{d}}{\mathrm{d}\theta}\big(d_1\cos^2\theta + d_2\sin^2\theta\big) \\ &= \sigma_\eta^2\Big[d_1 \cdot 2\cos\theta(-\sin\theta) + d_2 \cdot 2\sin\theta\cos\theta\Big] \\ &= 2\sigma_\eta^2 \sin\theta\cos\theta\,(d_2 - d_1).\end{aligned}$$

**Step 2: Quotient Expression.**

$$J'(\theta) \;=\; \frac{\mathcal{N}'(\theta)\,\mathcal{D}(\theta) - \mathcal{N}(\theta)\,\mathcal{D}'(\theta)}{\big[\mathcal{D}(\theta)\big]^2}.$$

Substitute expressions:

$$\begin{aligned}J'(\theta) &= \frac{\big[-2a^2\cos\theta\sin\theta\big]\,\big[\sigma_\eta^2(d_1\cos^2\theta + d_2\sin^2\theta)\big] \;-\; \big[a^2\cos^2\theta\big]\,\big[2\sigma_\eta^2\sin\theta\cos\theta(d_2 - d_1)\big]}{\big[\sigma_\eta^2(d_1\cos^2\theta + d_2\sin^2\theta)\big]^2} \\ &= \frac{-2a^2\sigma_\eta^2\cos\theta\sin\theta\,(d_1\cos^2\theta + d_2\sin^2\theta) \;+\; 2a^2\sigma_\eta^2\cos^3\theta\sin\theta\,(d_1 - d_2)}{\big[\sigma_\eta^2(d_1\cos^2\theta + d_2\sin^2\theta)\big]^2}.\end{aligned}$$

**Step 3: Simplification.** Factor $-2a^2\sigma_\eta^2\cos\theta\sin\theta$:

$$J'(\theta) = -\frac{2a^2\sigma_\eta^2\cos\theta\sin\theta\,\Big[d_1\cos^2\theta + d_2\sin^2\theta - (d_1 - d_2)\cos^2\theta\Big]}{\big[\sigma_\eta^2(d_1\cos^2\theta + d_2\sin^2\theta)\big]^2}.$$

It simplifies to:

$$d_1\cos^2\theta + d_2\sin^2\theta - d_1\cos^2\theta + d_2\cos^2\theta = d_2(\sin^2\theta + \cos^2\theta) = d_2.$$

Thus, we have in the end:

$$J'(\theta) = -\frac{2a^2\sigma_\eta^2\cos\theta\sin\theta\,d_2}{\big[\sigma_\eta^2(d_1\cos^2\theta + d_2\sin^2\theta)\big]^2}.$$

**Step 4: Zero condition.** Since $d_2 = (1 - \lambda_2)^{-2} > 0$ (by stability $|\lambda_2| < 1$), $a^2 > 0$ and $\sigma_\eta^2 > 0$ we have:

$$J'(\theta) = 0 \iff \cos\theta = 0 \;\text{ or }\; \sin\theta = 0.$$

Within $\theta \in \big[-\frac{\pi}{2}, \frac{\pi}{2}\big]$, stationary points are:

$$\theta \;\in\; \Big\{-\tfrac{\pi}{2},\, 0,\, \tfrac{\pi}{2}\Big\}.$$

#### B.4 Second Derivative

Following the first-derivative expression obtained,

$$J'(\theta) \;=\; -\,\frac{2\,a^2\,\sigma_\eta^2\;\cos\theta\;\sin\theta\;d_2}{\bigl[\sigma_\eta^2\bigl(d_1\cos^2\theta + d_2\sin^2\theta\bigr)\bigr]^2},$$

define

$$U(\theta) \equiv -\,2\,a^2\,\sigma_\eta^2\;d_2\;\cos\theta\sin\theta = -\,a^2\,\sigma_\eta^2\,d_2\;\sin(2\theta), \qquad V(\theta) \equiv \bigl[\sigma_\eta^2\bigl(d_1\cos^2\theta + d_2\sin^2\theta\bigr)\bigr]^2.$$

Then $J'(\theta) = U(\theta)/V(\theta)$ and

$$J''(\theta) \;=\; \frac{U'(\theta)\,V(\theta) - U(\theta)\,V'(\theta)}{V(\theta)^2}.$$

**Derivative of the Numerator.**

$$U'(\theta) \;=\; -\,2\,a^2\,\sigma_\eta^2\,d_2\,(\cos^2\theta - \sin^2\theta) = -\,2\,a^2\,\sigma_\eta^2\,d_2\;\cos(2\theta).$$

**Derivative of the Denominator.** Write $W(\theta) := d_1\cos^2\theta + d_2\sin^2\theta$, so $V(\theta) = \sigma_\eta^4 W(\theta)^2$.

$$W'(\theta) = \frac{\mathrm{d}}{\mathrm{d}\theta}\bigl(d_1\cos^2\theta + d_2\sin^2\theta\bigr) = -2d_1\sin\theta\cos\theta + 2d_2\sin\theta\cos\theta = -2(d_1 - d_2)\sin\theta\cos\theta.$$

Hence,

$$V'(\theta) = 2\,\sigma_\eta^4\,W(\theta)\,W'(\theta) = -\,4\,\sigma_\eta^4\,(d_1 - d_2)\,W(\theta)\;\cos\theta\sin\theta.$$

**at $\theta = 0$.**

$$U(0) = 0, \quad U'(0) = -\,2\,a^2\,\sigma_\eta^2\,d_2, \quad V(0) = \sigma_\eta^4 d_1^2, \quad V'(0) = 0.$$

Therefore,

$$J''(0) = \frac{U'(0)\,V(0) - U(0)\,V'(0)}{V(0)^2} = \frac{U'(0)}{V(0)}$$
$$= \frac{-2\,a^2\,\sigma_\eta^2\,d_2}{\sigma_\eta^4 d_1^2} = -\,\frac{2\,a^2\,d_2}{\sigma_\eta^2\,d_1^2} \;<\; 0.$$

If we set $\theta = 0$ so that we are at an extremum of $J(\theta)$ we see that $J''(0) < 0 \Rightarrow \theta = 0$ is a strict **local maximum**.

**The other two points.** Rather if we take $\theta = \pm\frac{\pi}{2}$, then since $\cos\bigl(\frac{\pi}{2}\bigr) = 0$ and $\sin\bigl(\frac{\pi}{2}\bigr) = 1$ the Fisher ratio itself vanishes:

$$J\Bigl(\pm\tfrac{\pi}{2}\Bigr) = \frac{a^2\cos^2\bigl(\frac{\pi}{2}\bigr)}{\sigma_\eta^2\bigl(d_1\cos^2\bigl(\frac{\pi}{2}\bigr) + d_2\sin^2\bigl(\frac{\pi}{2}\bigr)\bigr)} = \frac{0}{\sigma_\eta^2 d_2} = 0.$$

To classify the stationary point we do not need to compute the second-derivative here, because $J$ cannot be less than 0 (they have to be global minimums). Because $\theta \mapsto J(\theta)$ is even and $\pi$-periodic, these three points take up all stationary points; consequently, $\boldsymbol{v} = \pm\boldsymbol{u}_1$ uniquely maximizes the Fisher ratio, while $\boldsymbol{v} = \pm\boldsymbol{u}_\perp$ minimizes it.

#### B.5 Conclusion

The Fisher ratio is maximized only when the decoder axis is aligned to the axis of correlated variability:

$$\arg\max_{\|\boldsymbol{v}\|=1} J(\boldsymbol{v}) \;=\; \{\pm \boldsymbol{u}_1\}.$$

Therefore, once the stimulus response and axis of correlated variability are aligned, the best one-dimensional linear read-out axis is simply the axis of correlated variability itself.

### Appendix C – Single-axis Fisher information Changes

Here we ask a question: does setting the network's dominant noise mode to align with the stimulus response axis always improve the information available to a one–dimensional decoder? The answer depends on the constraints we assume for the decoder:

1. **No** *for an unconstrained 1-d decoder.* When the downstream read-out may choose **any** unit vector, the maximum single-axis Fisher information is fixed by the feed-forward drive $\mathbf{G}\boldsymbol{w}_F$ and the noise level $\sigma_\eta^2$; an optimal single-axis decoder can always whiten the noise, so its decoder direction is irrelevant.
2. **Yes** *for a mode-locked 1-d decoder.* When the read-out must coincide with a pre-selected network eigenmode, rotating that mode toward the stimulus axis, and optionally changing its eigenvalue, can raise the information.

#### C.1 Framework and notation recap

Recall that our network obeys:

##### Linear steady state

$$\boldsymbol{r} = (\mathbf{I} - \mathbf{A})^{-1}\big(\mathbf{G}\boldsymbol{w}_F\, s + \boldsymbol{\eta}\big), \qquad \boldsymbol{\eta} \sim \mathcal{N}(\mathbf{0}, \sigma_\eta^2 \mathbf{I}), \quad \rho(\mathbf{A}) < 1.$$

Throughout Appendix C we assume $\mathbf{A}$ is real-symmetric; thus an orthonormal eigen-basis exists: $\mathbf{A}\boldsymbol{u}_i = \lambda_i \boldsymbol{u}_i$ with $1 > \lambda_1 > \cdots > \lambda_N > -1$.

##### Stimulus Response and Noise Response statistics

$$\Delta\boldsymbol{\mu} = (\mathbf{I} - \mathbf{A})^{-1}\mathbf{G}\boldsymbol{w}_F = \sum_{i=1}^{N} \frac{c_i}{1-\lambda_i}\,\boldsymbol{u}_i, \quad c_i := \boldsymbol{u}_i^\top \mathbf{G}\boldsymbol{w}_F,$$

$$\Sigma = \sigma_\eta^2 (\mathbf{I} - \mathbf{A})^{-1}(\mathbf{I} - \mathbf{A})^{-\top} = \sigma_\eta^2 \sum_{i=1}^{N} d_i\, \boldsymbol{u}_i \boldsymbol{u}_i^\top, \quad d_i := \frac{1}{(1-\lambda_i)^2}.$$

##### Linear Fisher information for a unit decoder $\boldsymbol{v}$

$$J(\boldsymbol{v}) = \frac{(\boldsymbol{v}^\top \Delta\boldsymbol{\mu})^2}{\boldsymbol{v}^\top \Sigma \boldsymbol{v}}.$$

#### C.2 If the 1-d decoder may choose any direction

**Proposition 6** (Whitening). *The unit vector* $\boldsymbol{v}_{opt} = \dfrac{\Sigma^{-1}\Delta\boldsymbol{\mu}}{\|\Sigma^{-1}\Delta\boldsymbol{\mu}\|}$ *maximizes* $J(\boldsymbol{v})$*. The maximum is:*

$$J_{\max} = \Delta\boldsymbol{\mu}^\top \Sigma^{-1} \Delta\boldsymbol{\mu} = \frac{1}{\sigma_\eta^2} \sum_i \frac{\big(c_i/(1-\lambda_i)\big)^2}{d_i} = \frac{1}{\sigma_\eta^2} \sum_i c_i^2.$$

**Proof:** Since $J(\alpha\boldsymbol{v}) = J(\boldsymbol{v})$ for all $\alpha \neq 0$, we may restrict to $\|\boldsymbol{v}\| = 1$ without loss of generality. Let $\Sigma^{1/2}$ be the positive-definite square root and define whitened variables:

$$\widetilde{\boldsymbol{v}} := \Sigma^{1/2}\boldsymbol{v}, \qquad \widetilde{\Delta\boldsymbol{\mu}} := \Sigma^{-1/2}\Delta\boldsymbol{\mu}.$$

This yields: $\boldsymbol{v}^\top \Sigma \boldsymbol{v} = \widetilde{\boldsymbol{v}}^\top \widetilde{\boldsymbol{v}}$ and $\boldsymbol{v}^\top \Delta\boldsymbol{\mu} = \widetilde{\boldsymbol{v}}^\top \widetilde{\Delta\boldsymbol{\mu}}$. Thus

$$J(\boldsymbol{v}) = \frac{(\widetilde{\boldsymbol{v}}^\top \widetilde{\Delta\boldsymbol{\mu}})^2}{\widetilde{\boldsymbol{v}}^\top \widetilde{\boldsymbol{v}}} \quad \text{for} \quad \widetilde{\boldsymbol{v}} \neq \mathbf{0}.$$

The Cauchy-Schwarz inequality gives $(x^\top y)^2 \leq (x^\top x)(y^\top y)$ with equality iff $x \parallel y$. Substituting $x = \widetilde{\boldsymbol{v}}$, $y = \widetilde{\Delta\boldsymbol{\mu}}$ gives

$$J(\boldsymbol{v}) \;\leq\; \widetilde{\Delta\boldsymbol{\mu}}^\top \widetilde{\Delta\boldsymbol{\mu}} = \Delta\boldsymbol{\mu}^\top \Sigma^{-1} \Delta\boldsymbol{\mu},$$

with equality when $\Sigma^{1/2}\boldsymbol{v} \propto \Sigma^{-1/2}\Delta\boldsymbol{\mu}$.

The maximizer (up to sign) is therefore

$$\boldsymbol{v}_{\text{opt}} \propto \Sigma^{-1}\Delta\boldsymbol{\mu}, \qquad \boldsymbol{v}_{\text{opt}} = \frac{\Sigma^{-1}\Delta\boldsymbol{\mu}}{\|\Sigma^{-1}\Delta\boldsymbol{\mu}\|},$$

and the maximal information is

$$J_{\max} = \Delta\boldsymbol{\mu}^\top \Sigma^{-1} \Delta\boldsymbol{\mu} = \frac{1}{\sigma_\eta^2} \sum_i \frac{(c_i/(1-\lambda_i))^2}{d_i} = \sum_i \frac{c_i^2}{\sigma_\eta^2},$$

since $d_i = 1/(1-\lambda_i)^2$. □

The identity $\sum_{i=1}^N c_i^2 = \|\mathbf{G}\boldsymbol{w}_F\|^2$ reveals: $J_{\max} = \|\mathbf{G}\boldsymbol{w}_F\|^2/\sigma_\eta^2$ is fixed for given inputs and noise. As a result modifications to the recurrent matrix $\mathbf{A}$ cannot affect $J_{\max}$.

#### C.3 Decoder locked to a single eigen-axis

Assume the downstream circuit can only read out a single network eigenmode, say the $k$-th eigenvector $\boldsymbol{u}_k$. Hence the weight vector is fixed to

$$\boldsymbol{v} = \pm \boldsymbol{u}_k.$$

With angle $\vartheta_k := \angle(\Delta\boldsymbol{\mu}, \boldsymbol{u}_k)$ the locked-axis Fisher information equals

$$J_{\mathbf{before}} = \frac{(\boldsymbol{u}_k^\top \Delta\boldsymbol{\mu})^2}{\sigma_\eta^2 d_k} = \underbrace{\frac{\|\Delta\boldsymbol{\mu}\|^2}{\sigma_\eta^2 d_k}}_{J_{\max} \text{ on this axis}} \times \cos^2\vartheta_k.$$

When $\vartheta_k > 0$ we lose the factor $\cos^2\vartheta_k$ compared with perfect alignment.

We examine two operations on $\boldsymbol{u}_k$:
**A.** Pure rotation: turn $\boldsymbol{u}_k$ without changing its eigenvalue $\lambda_k$.
**B.** Rotation *plus* a change of $\lambda_k$ (making that mode slower or faster).

##### Operation A – pure rotation (eigenvalue unchanged to first order)

We rotate the decoder axis from the $k$-th eigenvector $\boldsymbol{u}_k$ toward the stimulus-response vector

$$\widehat{\Delta\boldsymbol{\mu}} := \frac{\Delta\boldsymbol{\mu}}{\|\Delta\boldsymbol{\mu}\|}, \qquad \vartheta_k := \angle\big(\Delta\boldsymbol{\mu}, \boldsymbol{u}_k\big) \in (0, \tfrac{\pi}{2}],$$

while leaving the eigenvalue $\lambda_k$ invariant up to $\mathcal{O}(\vartheta_k^2)$.

**Perturbation.** Set

$$\delta\mathbf{A} \;=\; \varepsilon\,\big(\boldsymbol{u}_k\boldsymbol{w}^\top + \boldsymbol{w}\boldsymbol{u}_k^\top\big), \qquad \varepsilon := \tan\vartheta_k, \tag{28}$$

where

$$\boldsymbol{w} := \widehat{\Delta\boldsymbol{\mu}} \;-\; \big(\boldsymbol{u}_k^\top\widehat{\Delta\boldsymbol{\mu}}\big)\,\boldsymbol{u}_k \quad (\;\boldsymbol{w}\perp\boldsymbol{u}_k\;).$$

*Symmetry:* $\delta\mathbf{A}$ is symmetric by construction; hence the perturbed matrix $\mathbf{A}+\delta\mathbf{A}$ remains diagonalizable with real eigenvalues.

**No first-order eigenvalue shift.** Because $\boldsymbol{w}\perp\boldsymbol{u}_k$,

$$\boldsymbol{u}_k^\top\delta\mathbf{A}\,\boldsymbol{u}_k \;=\; \varepsilon\;\underbrace{\boldsymbol{u}_k^\top\boldsymbol{u}_k}_{=1}\;\underbrace{\boldsymbol{w}^\top\boldsymbol{u}_k}_{=0}+\varepsilon\;\underbrace{\boldsymbol{u}_k^\top\boldsymbol{w}}_{=0}\;\underbrace{\boldsymbol{u}_k^\top\boldsymbol{u}_k}_{=1}=0,$$

so the first-order eigenvalue correction is $\delta\lambda_k = 0\;+\mathcal{O}(\varepsilon^2)$.

**First-order eigenvector correction.** For a non-degenerate eigenvalue of a symmetric matrix

$$\delta\boldsymbol{u}_k \;=\; \sum_{j\neq k}\frac{\boldsymbol{u}_j^\top\delta\mathbf{A}\,\boldsymbol{u}_k}{\lambda_k-\lambda_j}\,\boldsymbol{u}_j = \varepsilon\sum_{j\neq k}\frac{\boldsymbol{u}_j^\top\boldsymbol{w}}{\lambda_k-\lambda_j}\,\boldsymbol{u}_j = \mathcal{O}(\varepsilon)\,\boldsymbol{w} \quad\propto\quad \boldsymbol{w}.$$

Consequently $\boldsymbol{u}_k+\delta\boldsymbol{u}_k = \widehat{\Delta\boldsymbol{\mu}}+\mathcal{O}(\varepsilon^2)$, i.e. the decoder axis is rotated exactly by $\vartheta_k$ to first order.

**Effect on single-axis Fisher information.** Decoder variance remains $\boldsymbol{u}_k^\top\Sigma\boldsymbol{u}_k=\sigma_\eta^2 d_k$ because $d_k$ did not change at $\mathcal{O}(\varepsilon)$. The signal projection increases from $\|\Delta\boldsymbol{\mu}\|\cos\vartheta_k$ to $\|\Delta\boldsymbol{\mu}\|$. Therefore

$$J_{\textbf{after}}=\frac{\|\Delta\boldsymbol{\mu}\|^2}{\sigma_\eta^2 d_k}=\frac{J_{\textbf{before}}}{\cos^2\vartheta_k}, \qquad J_{\textbf{after}}>J_{\textbf{before}} \;\text{ for } 0<\vartheta_k<\tfrac{\pi}{2}.$$

A pure rotation that keeps the eigenvalue fixed to first order *always* increases the single-axis Fisher information.

##### Operation B – rotation *and* eigenvalue change

Now let the rotation also modify the eigenvalue:

$$\lambda_k^{\textbf{before}} \;\longrightarrow\; \lambda_k^{\textbf{after}}, \quad d_k^{\textbf{after}} = d_k^{\textbf{before}}\,R_d, \quad R_d := \Big(\frac{1-\lambda_k^{\textbf{before}}}{1-\lambda_k^{\textbf{after}}}\Big)^2 > 1.$$

Assume the same perturbation scales the signal norm by $g := \|\Delta\boldsymbol{\mu}_{\textbf{after}}\|/\|\Delta\boldsymbol{\mu}_{\textbf{before}}\|$ (and the angle is now zero).

Changing only the $k$-th eigenvalue $\lambda_k^{\text{before}}\to\lambda_k^{\text{after}}$ leaves the drive coefficient $c_k=\boldsymbol{u}_k^\top\mathbf{G}\boldsymbol{w}_F$ unchanged, so we have:

$$\Delta\mu_k^{\text{before}}=\frac{c_k}{1-\lambda_k^{\text{before}}}, \qquad \Delta\mu_k^{\text{after}}=\frac{c_k}{1-\lambda_k^{\text{after}}},$$

$$d_k^{\text{before}}=\frac{\sigma_\eta^2}{(1-\lambda_k^{\text{before}})^2}, \qquad d_k^{\text{after}}=\frac{\sigma_\eta^2}{(1-\lambda_k^{\text{after}})^2}.$$

Hence the signal-gain and variance-gain factors are:

$$g \;=\; \frac{\|\Delta\boldsymbol{\mu}_{\text{after}}\|}{\|\Delta\boldsymbol{\mu}_{\text{before}}\|}=\frac{1-\lambda_k^{\text{before}}}{1-\lambda_k^{\text{after}}}, \qquad R_d \;=\; \frac{d_k^{\text{after}}}{d_k^{\text{before}}}=\Big(\frac{1-\lambda_k^{\text{before}}}{1-\lambda_k^{\text{after}}}\Big)^2.$$

**New information.**

$$J_{\textbf{after}} = \frac{g^2 \|\Delta \boldsymbol{\mu}_{\textbf{before}}\|^2}{\sigma_\eta^2 d_k^{\textbf{before}} R_d} = J_{\textbf{before}} \, \frac{g^2}{R_d}.$$

**Gain condition.**

$$J_{\textbf{after}} > J_{\textbf{before}} \quad \Longleftrightarrow \quad g > \sqrt{R_d}.$$

Rotating onto a *slower / noisier* axis helps only if the network provides *extra* mean gain that counters the $\sqrt{\text{variance}}$ penalty. In a purely linear symmetric network mean and variance grow in lock-step ($g^2 = R_d$), so the inequality is never satisfied; some selective or nonlinear amplification must be present.

### Appendix D – Optimal encoding for temporal discrimination

Here we revisit the encoding question in a setting that explicitly models *temporal integration*. We show that, when the population evolves over time and a downstream unit reads out the state at a finite decision time $T$, the encoding that maximises discriminability $d'$—even with an *unconstrained* decoder—concentrates all stimulus power on the *slowest* recurrent mode. Because the slowest mode is also the dominant noise direction, this provides a normative, encoding-side rationale for the alignment between the stimulus axis and the axis of correlated variability observed in our data.

We proceed in four steps:

1. Set up the temporal dynamics and project onto the eigenbasis (D.1).
2. Compute signal and noise at decision time $T$ for each mode (D.2).
3. For a *fixed* encoding, derive the best one-dimensional decoder and the resulting $d'^2$ (D.3).
4. Optimise over encodings to show that all stimulus power should go on the slowest mode (D.4–D.5).

#### D.1 Setup

**Noise model.** Appendices A–C analyse a steady-state snapshot model in which each neuron receives a single trial-to-trial noise draw $\boldsymbol{\eta} \sim \mathcal{N}(\mathbf{0}, \sigma_\eta^2 \mathbf{I})$, which is then filtered by the recurrence. To study temporal integration, we instead model within-trial fluctuations by replacing $\boldsymbol{\eta}$ with a temporally white noise process $\sigma_\eta \, \boldsymbol{\xi}(t)$, where $\boldsymbol{\xi}(t) \in \mathbb{R}^N$ is a vector of independent standard white-noise processes.

**Dynamics and eigenbasis.** We take Eq. (1) with $\mathbf{G} = \mathbf{I}$, $\tau = 1$, and $\mathbf{W}_\mathrm{R}$ real-symmetric:

$$\dot{\boldsymbol{r}}(t) = -\boldsymbol{r}(t) + \mathbf{W}_\mathrm{R} \, \boldsymbol{r}(t) + \boldsymbol{w}_\mathrm{F} \, s + \sigma_\eta \, \boldsymbol{\xi}(t), \tag{29}$$

where $\xi_i(t)$ denotes the $i$-th component of $\boldsymbol{\xi}(t)$ and satisfies $\langle \xi_i(t) \, \xi_j(t') \rangle = \delta_{ij} \, \delta(t - t')$ (here $\delta_{ij}$ is the Kronecker delta and $\delta(\cdot)$ is the delta function). Let $\mathbf{W}_\mathrm{R} \boldsymbol{u}_i = \lambda_i \boldsymbol{u}_i$ with $1 > \lambda_1 > \lambda_2 \geq \cdots \geq \lambda_N > -1$, and define the decay rate of each mode as $\gamma_i \coloneqq 1 - \lambda_i > 0$; thus $\gamma_1 < \gamma_2 \leq \cdots \leq \gamma_N$ and mode 1 is the slowest.

**Task and encoding.** A constant stimulus $s \in \{+\frac{\Delta s}{2}, -\frac{\Delta s}{2}\}$ is presented over $[0, T]$. Projecting (29) onto the eigenbasis by defining $\alpha_i(t) \coloneqq \boldsymbol{u}_i^\top \boldsymbol{r}(t)$ (the activity of mode $i$) gives $N$ independent Ornstein–Uhlenbeck processes:

$$\dot{\alpha}_i(t) = -\gamma_i \, \alpha_i(t) + f_i \, s + \sigma_\eta \, \xi_i(t), \qquad \alpha_i(0) = 0, \tag{30}$$

where $f_i \coloneqq \boldsymbol{u}_i^\top \boldsymbol{w}_\mathrm{F}$ is the projection of the feedforward drive onto mode $i$; the initial condition $\alpha_i(0) = 0$ corresponds to stimulus onset from a baseline state. The *encoding* is the choice of $\boldsymbol{f} = (f_1, \ldots, f_N)$ subject to a total stimulus-power budget $\sum_i f_i^2 = P$. Our goal is to find the encoding that maximises the discriminability $d'$ available to the best one-dimensional decoder at time $T$.

#### D.2 Signal and noise at decision time $T$

Each mode in (30) is a scalar OU process with known solution. At time $T$, the conditional mean and variance are:

**Signal.**

$$\Delta\mu_i(T) := \mathbb{E}[\alpha_i(T)\,|\,s{=}\tfrac{\Delta s}{2}] - \mathbb{E}[\alpha_i(T)\,|\,s{=}-\tfrac{\Delta s}{2}] = f_i\,\Delta s\;G_i(T), \qquad G_i(T) := \frac{1-e^{-\gamma_i T}}{\gamma_i}, \tag{31}$$

where $G_i(T)$ is the temporal gain of mode $i$: the integrated effect of a constant drive filtered through an exponential decay. For $T \ll 1/\gamma_i$ the signal grows approximately linearly, $G_i(T) \approx T$; at steady state $G_i \to 1/\gamma_i$. Slow modes (small $\gamma_i$) therefore accumulate a larger integrated signal.

**Noise.**

$$V_i(T) := \mathrm{Var}[\alpha_i(T)] = \sigma_\eta^2\,\frac{1-e^{-2\gamma_i T}}{2\gamma_i}\,, \qquad \mathrm{cov}\big(\alpha_i(T), \alpha_j(T)\big) = 0 \;\; (i \neq j). \tag{32}$$

For $T \ll 1/\gamma_i$ the variance grows as $V_i(T) \approx \sigma_\eta^2 T$ (random-walk regime); at steady state $V_i \to \sigma_\eta^2/(2\gamma_i)$. Moreover, for any fixed $T > 0$ the function $V(\gamma, T) = \sigma_\eta^2(1 - e^{-2\gamma T})/(2\gamma)$ is strictly decreasing in $\gamma$, so $V_1(T) > V_2(T) \geq \cdots \geq V_N(T)$: the slowest mode carries the most noise at every decision time.

Under the snapshot noise model of A–C the steady-state variance in mode $i$ scales as $1/\gamma_i^2$, because the recurrence filters a single draw rather than integrating diffusion. The steady-state mean response $\boldsymbol{\Delta\mu}$ (as $T \to \infty$) is identical in both formulations; only the noise accumulation differs.

#### D.3 Best one-dimensional decoder for a fixed encoding

*In this section the encoding* $\boldsymbol{f}$ *is held fixed; we optimise over encodings in D.5.*

A downstream unit computes $y = \sum_i w_i\,\alpha_i(T)$ with weights $\boldsymbol{w}$. Because the modes are independent at time $T$, the discriminability is

$$d'(\boldsymbol{w}) = \frac{\big|\sum_i w_i\,\Delta\mu_i(T)\big|}{\sqrt{\sum_i w_i^2\,V_i(T)}} = \frac{\big|\Delta s \sum_i w_i f_i G_i(T)\big|}{\sqrt{\sum_i w_i^2\,V_i(T)}}. \tag{33}$$

To maximise (33) over $\boldsymbol{w}$, define $a_i := f_i\,G_i(T)$ (signal coefficient) and $b_i := V_i(T)$ (noise variance). Then

$$d'^{\,2}(\boldsymbol{w}) = \Delta s^2\,\frac{\big(\sum_i w_i a_i\big)^2}{\sum_i w_i^2 b_i}.$$

Applying the Cauchy–Schwarz inequality with $u_i := w_i\sqrt{b_i}$ and $v_i := a_i/\sqrt{b_i}$,

$$\Big(\sum_i w_i a_i\Big)^2 = \Big(\sum_i u_i\,v_i\Big)^2 \leq \Big(\sum_i u_i^2\Big)\Big(\sum_i v_i^2\Big) = \Big(\sum_i w_i^2 b_i\Big)\Big(\sum_i \frac{a_i^2}{b_i}\Big),$$

with equality when $w_i^* \propto a_i/b_i = f_i\,G_i(T)/V_i(T)$. Substituting back:

$$d'^{\,2}_{\mathrm{opt}} = \Delta s^2 \sum_{i=1}^{N} \frac{f_i^2\,G_i(T)^2}{V_i(T)}. \tag{34}$$

We now simplify the summand. Using the factorisation $1 - e^{-2\gamma_i T} = (1 - e^{-\gamma_i T})(1 + e^{-\gamma_i T})$,

$$\frac{G_i(T)^2}{V_i(T)} = \frac{(1-e^{-\gamma_i T})^2/\gamma_i^2}{\sigma_\eta^2(1-e^{-\gamma_i T})(1+e^{-\gamma_i T})/(2\gamma_i)} = \frac{1}{\sigma_\eta^2}\,\frac{2(1-e^{-\gamma_i T})}{\gamma_i(1+e^{-\gamma_i T})}.$$

Define the noise-amplitude-independent weight

$$\widetilde{H}_i(T) := \sigma_\eta^2\,\frac{G_i(T)^2}{V_i(T)} = \frac{2(1-e^{-\gamma_i T})}{\gamma_i(1+e^{-\gamma_i T})} = \frac{2}{\gamma_i}\,\tanh\!\Big(\frac{\gamma_i T}{2}\Big), \tag{35}$$

so that (34) becomes

$$\boxed{d'^2_{\text{opt}} = \frac{\Delta s^2}{\sigma_\eta^2} \sum_{i=1}^{N} f_i^2\, \widetilde{H}_i(T).} \tag{36}$$

This is the best discriminability achievable by *any* one-dimensional decoder for a given encoding $\boldsymbol{f}$. The encoding question reduces to: *which allocation of* $\{f_i^2\}$ *under* $\sum_i f_i^2 = P$ *maximises* (36)*?*

#### D.4 $\widetilde{H}(\gamma, T)$ decreases with $\gamma$

**Lemma 1.** *For fixed* $T > 0$*, the map* $\gamma \mapsto \widetilde{H}(\gamma, T)$ *in* (35) *is strictly decreasing on* $(0, \infty)$*.*

*Proof.* Substitute $u \coloneqq \gamma T > 0$:

$$\widetilde{H}(\gamma, T) = \frac{2T}{u} \tanh\!\Big(\frac{u}{2}\Big).$$

It suffices to show $\varphi(u) \coloneqq \tanh(u/2)/u$ is strictly decreasing. Differentiate:

$$\varphi'(u) = \frac{u\, \operatorname{sech}^2(u/2)/2 - \tanh(u/2)}{u^2} = \frac{u - \sinh u}{2u^2 \cosh^2(u/2)} < 0,$$

since $\sinh u > u$ for all $u > 0$. □

Because $\gamma_1 < \gamma_2 \le \cdots \le \gamma_N$, the lemma gives:

$$\widetilde{H}_1(T) > \widetilde{H}_2(T) \ge \cdots \ge \widetilde{H}_N(T). \tag{37}$$

#### D.5 Optimal encoding

**Proposition 7.** *Given decay rates* $0 < \gamma_1 < \gamma_2 \le \cdots \le \gamma_N$ *and a stimulus-power budget* $\sum_i f_i^2 = P$*, the encoding that maximises* $d'^2_{\text{opt}}$ *in* (36) *is*

$$f_i^* = \pm\sqrt{P}\, \delta_{i,1},$$

*i.e. all stimulus power is placed on the slowest mode.*

*Proof.* Write $p_i \coloneqq f_i^2 \ge 0$ with $\sum_i p_i = P$. The objective $\sum_i p_i\, \widetilde{H}_i(T)$ is linear in $\boldsymbol{p}$ over the simplex $\{p_i \ge 0 : \sum_i p_i = P\}$. A linear function over a simplex is maximised at a vertex. By (37) the vertex with the largest objective is $p_1 = P$, $p_i = 0$ for $i > 1$. □

#### D.6 Connection to information-limiting correlations

Moreno-Bote et al. (2014) showed that correlations proportional to $\boldsymbol{f}_\mu \boldsymbol{f}_\mu^\top$ (where $\boldsymbol{f}_\mu \equiv \partial \boldsymbol{\mu}/\partial s$) limit snapshot Fisher information. In our linear recurrent network the dominant noise direction is also the slowest mode, and Proposition 7 shows that temporal integration makes it optimal to place stimulus power on that same slow mode despite its larger correlated variability.

Together with Appendix B, this establishes the encoding–decoding logic: (i) optimal encoding places stimulus power on the slowest recurrent mode to exploit temporal integration (this appendix D); (ii) once the stimulus and noise axes are aligned, the optimal one-dimensional readout lies along the same direction (Appendix B).
